# An inhibitory circuit-based enhancer of Dyrk1a function reverses *Dyrk1a*-associated impairment in social recognition

**DOI:** 10.1101/2023.02.03.526955

**Authors:** Yu-Tzu Shih, Jason Bondoc Alipio, Amar Sahay

## Abstract

Heterozygous mutations in the Dual specificity tyrosine-phosphorylation-regulated kinase 1a *Dyrk1a* gene define a syndromic form of Autism Spectrum Disorder. The synaptic and circuit mechanisms mediating Dyrk1a functions in social cognition are unclear. Here, we identify a social experience-sensitive mechanism in hippocampal mossy fiber-parvalbumin interneuron (PV IN) synapses by which Dyrk1a recruits feedforward inhibition of CA3 and CA2 to promote social recognition. We employ genetic epistasis logic to identify a cytoskeletal protein, Ablim3, as a synaptic substrate of Dyrk1a. We demonstrate that *Ablim3* downregulation in dentate granule cells of adult hemizygous *Dyrk1a* mice is sufficient to restore PV IN mediated inhibition of CA3 and CA2 and social recognition. Acute chemogenetic activation of PV INs in CA3/CA2 of adult hemizygous *Dyrk1a* mice also rescued social recognition. Together, these findings illustrate how targeting Dyrk1a synaptic and circuit substrates as “enhancers of Dyrk1a function” harbors potential to reverse *Dyrk1a* haploinsufficiency-associated circuit and cognition impairments.

**Highlights:** Dyrk1a in mossy fibers recruits PV IN mediated feed-forward inhibition of CA3 and CA2

Dyrk1a-Ablim3 signaling in mossy fiber-PV IN synapses promotes inhibition of CA3 and CA2

Downregulating *Ablim3* restores PV IN excitability, CA3/CA2 inhibition and social recognition in *Dyrk1a+/-* mice

Chemogenetic activation of PV INs in CA3/CA2 rescues social recognition in *Dyrk1a+/-* mice

## INTRODUCTION

Autism spectrum disorder (ASD) is a class of heterogeneous neurodevelopmental disorders that affects 1 in 100 children around the world and is characterized by impaired social cognition, repetitive behaviors and increased seizure risk (Lord et al., 2020). A fundamental challenge in identifying new treatments for ASD is understanding how disease gene mutations, grounded in robust human genetic signals, affect cellular, synaptic and circuit mechanisms underlying behavior (Willsey et al., 2022). Human genetic studies and large-scale exome-sequencing efforts implicate haploinsufficient loss-of-function (LOF) mutations in the Dual specificity tyrosine-phosphorylation-regulated kinase1a *Dyrk1a* gene in syndromic ASD (Earl et al., 2017; O’Roak et al., 2012; Satterstrom et al., 2020). However, the synaptic and circuit mechanisms by which Dyrk1a affects social cognition are poorly understood. Currently, there are no drugs that enhance or restore Dyrk1a function. This challenge is magnified because *Dyrk1a* is ubiquitously expressed in the developing and mature brain and Dyrk1a phosphorylates more than 30 substrates that regulate diverse processes such as cell-cycle, neuronal migration, axonal and dendritic development, synapse formation, synaptic signaling and function (Arranz et al., 2019; Atas-Ozcan et al., 2021; Benavides-Piccione et al., 2005; Chen et al., 2014; Classen et al., 2020; Dang et al., 2018; Duchon and Herault, 2016; Guedj et al., 2012; Levy et al., 2021; Soppa and Becker, 2015; Souchet et al., 2014; Thompson et al., 2015; Vidaki et al., 2017).

One approach to developing strategies to restore Dyrk1a function is to assess the potential for targeting Dyrk1a synaptic substrates in social cognition circuits to reverse circuit- and social cognition impairments associated with *Dyrk1a* hemizygosity. Genetic epistasis guided logic (Avery and Wasserman, 1992; Domingo et al., 2019) posits that potentiation of a downstream effector (i.e. synaptic substrate) of Dyrk1a function should rescue Dyrk1a dependent synaptic impairments and associated neural-circuit functions. Furthermore, although *Dyrk1a* haploinsufficiency associated impairments in social cognition arise from reduced Dyrk1a mediated signaling and recruitment of different substrates throughout development, it is plausible that a subset of synaptic substrates may be targeted in adulthood to alleviate cognitive impairments.

Dentate gyrus (DG)-CA3/CA2 circuits play a critical role in encoding social experiences by integrating sensory information from association cortices, entorhinal cortex, prefrontal cortex and subcortical sites (Alexander et al., 2018; Alexander et al., 2016; Chen et al., 2020; Cope et al., 2022; Finlay et al., 2015; Gangopadhyay et al., 2021; Hitti and Siegelbaum, 2014; Lehr et al., 2021; Lin et al., 2018; Lopez-Rojas et al., 2022; Meira et al., 2018; Montagrin et al., 2018; Oliva et al., 2016; Oliva et al., 2020; Raam et al., 2017; Schafer and Schiller, 2018a, b; Stevenson and Caldwell, 2014; Tavares et al., 2015; Tuncdemir et al., 2022; Woods et al., 2020). Prior work has shown that experience or learning increases dentate granule cell (DGC) recruitment of parvalbumin inhibitory neurons (PV IN) mediated perisomatic inhibition of CA3 neurons (Guo et al., 2018). Anatomically, this manifests as an increase in number of mossy fiber terminal (MFT) filopodial contacts with PV INs and increase in number of PV IN perisomatic contacts in CA3 (Acsady et al., 1998; Caroni, 2015; Guo et al., 2018; Ruediger et al., 2012; Ruediger et al., 2011; Twarkowski et al., 2022). PV mediated perisomatic inhibitory synaptic transmission in downstream principal neurons (CA3/CA2) may facilitate encoding, routing and consolidation of experiential information in distinct neuronal ensembles by imposing a sparse activity regimen (Mori et al., 2007; Neubrandt et al., 2017; Pelkey et al., 2017; Torborg et al., 2010), synchronizing activity of principal cells and generation of network oscillations (Amilhon et al., 2015; Buzsaki, 2015; Cardin et al., 2009; Csicsvari et al., 2000; de la Prida et al., 2006; Gan et al., 2016; Hu et al., 2014; Klausberger et al., 2003; Ognjanovski et al., 2017; Sohal et al., 2009). Consistently, we recently demonstrated that increasing feed-forward inhibition (FFI) in DG-CA3 resulted in enhanced stability and precision of context-associated neuronal ensembles and hippocampal-cortical synchrony (Twarkowski et al., 2022). Whether DGC recruitment of PV IN mediated inhibition of CA3 and CA2 is necessary for encoding social experiences is not known.

Studies using non-neuronal cells have shown that Dyrk1a phosphorylates and functionally inactivates a family of F-actin stabilizing proteins, Ablim 1-3, that results in destabilization of branched F-actin (De Toma et al., 2019; Schneider et al., 2015; Singh and Lauth, 2017). This was interesting to us as we had previously shown that *Ablim3* downregulation in DGCs results in generation of MFT-filopodial contacts with PV INs, a process that is thought to depend on destabilization of branched F-actin in MFTs (Guo et al., 2018). Based on these observations, we hypothesized that Dyrk1a recruits Ablim3 in mossy fibers to promote DGC recruitment of PV IN mediated GABAergic inhibition in CA3 and CA2 to support social cognition. Furthermore, using functional genetic epistasis logic, we reasoned that if Ablim3 functions downstream of Dyrk1a in mossy fiber terminals (MFTs), then targeting *Ablim3* in DGCs may rescue *Dyrk1a* LOF associated impairments in social cognition.

Here, we show that social experience increases MFT-filopodial contacts with PV INs and PV IN perisomatic contacts in CA3 and CA2 pyramidal neurons. We demonstrate that *Dyrk1a* is necessary in DGCs of adult mice for feed-forward inhibition connectivity (MFT-filopodial contacts with PV INs), PV IN perisomatic contacts in CA3/CA2 and social recognition. We also demonstrate that Ablim3 functions downstream of Dyrk1a or is epistatic to Dyrk1a in mossy fibers. Based on these data, we use hemizygous *Dyrk1a* mice to show that loss of one *Dyrk1a* allele throughout development resulted in reduced MFT-filopodial contacts with PV INs, PV IN intrinsic excitability, PV IN perisomatic contacts in CA3/CA2 neurons and impaired social recognition in adulthood. These anatomical alterations were mirrored by reduced mossy fiber excitatory drive onto PV INs in CA3 and CA2 and mossy fiber recruitment of GABAergic inhibition of CA3 and CA2. Viral downregulation of *Ablim3* in DGCs of adult hemizygous *Dyrk1a* mice reversed these developmental anatomical and electrophysiological deficits and restored social recognition. Motivated by these findings, we assessed the impact of selectively activating PV INs in CA3/CA2 of adult hemizygous *Dyrk1a* mice on social recognition. We found that acute chemogenetic activation of PV INs in CA3/CA2 was sufficient to rescue social recognition impairments. Together, these findings illuminate how targeting Dyrk1a synaptic substrates as “circuit-based enhancers of Dyrk1a function” may harbor potential to reverse *Dyrk1a* haploinsufficiency-associated circuit and cognition impairments.

## RESULTS

### *Dyrk1a* is required in DGCs of adult mice for FFI connectivity, PV IN perisomatic contacts and social recognition

Learning increases mossy fiber excitatory drive onto PV INs (Caroni, 2015; Guo et al., 2018; Ruediger et al., 2012; Ruediger et al., 2011; Twarkowski et al., 2022) and enhances PV IN perisomatic contacts (“PV IN plasticity”) in CA3 (Guo et al., 2018; Twarkowski et al., 2022). To determine if social experience also increases FFI connectivity and PV IN plasticity in CA3/CA2, lentiviruses expressing GFP (CaMKIIα-GFP) were injected into the DG of 2-months old *Dyrk1a^+/+^* mice two weeks prior to habituation to a context over 7 days. On day 8, mice were exposed to either the same context (context) or a novel juvenile mouse (social stimulus, same sex and strain) on day 8 **(Figure 1A)**. 1.5 hours following social or context exposure we analyzed vGLUT1+ (vesicular glutamate transporter 1) mossy fiber terminal (MFT) filopodial contacts with PV INs and PV IN contacts (puncta) in CA3 and CA2 by immunohistochemistry. Social experience increased MFT-filopodial contacts with PV INs and PV IN contacts or “PV IN puncta density” in CA3 and CA2 **(Figures 1B-1C)**.

**Figure 1.**
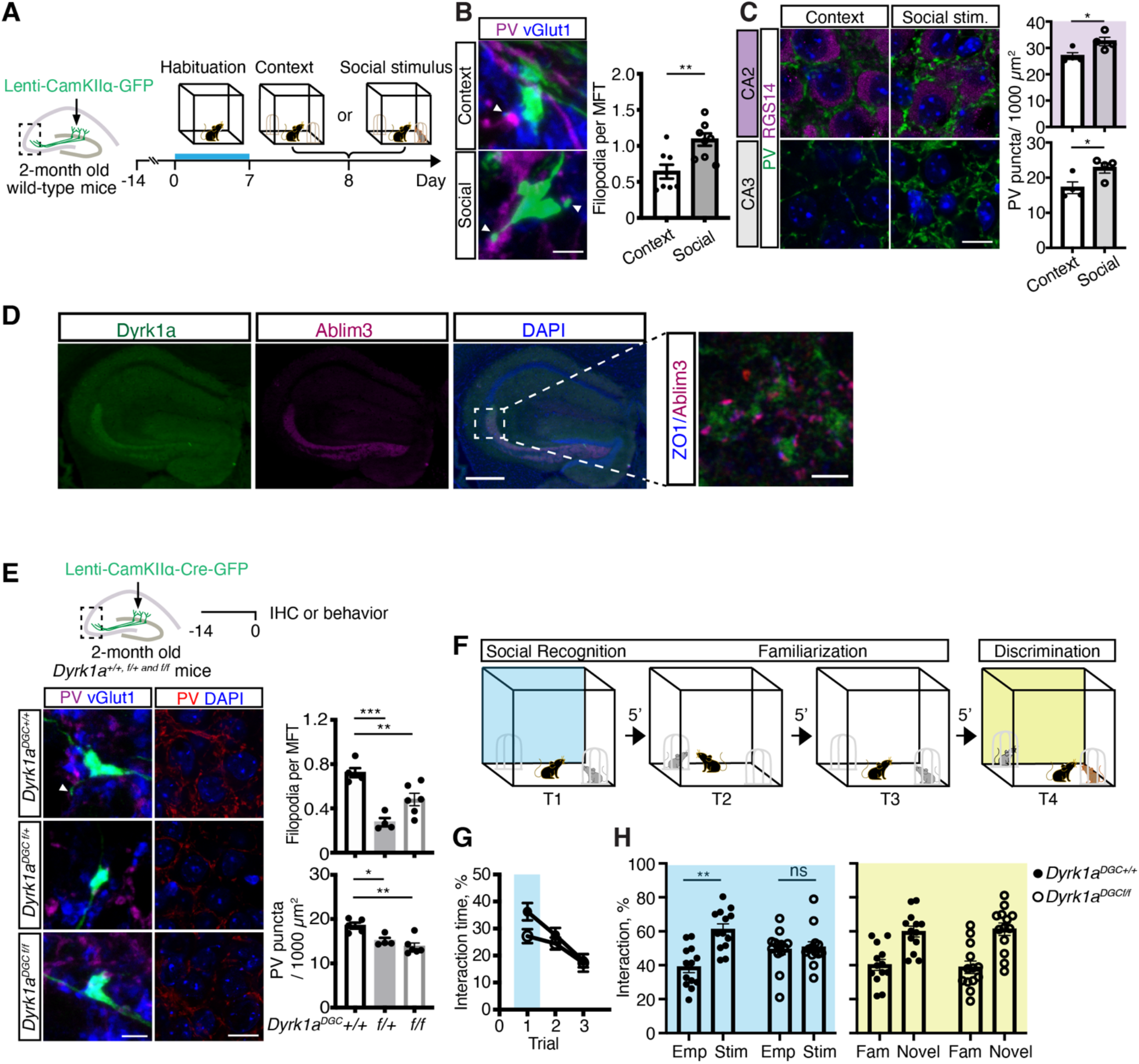
*Dyrk1a* is required in DGCs of adult mice for FFI connectivity, PV IN perisomatic contacts and social recognition. (A) Experimental design. Lentiviruses expressing GFP (CaMKIIα-GFP) were injected into the DG of 2-month old *Dyrk1a^+/+^* mice 2 weeks prior to habituation to a chamber (Context) for 7 days. On day 8 mice were exposed to the familiarized context or familiarized context with a social stimulus (novel juvenile mouse). (B) Representative images showing GFP^+^ MFTs (filopodial extensions, arrows) and quantification of the number of filopodia per vGLUT1^+^ MFTs (N = 8 mice per group) in CA3. Scale bar, 5 μm. p < 0.01 using twotailed unpaired *t* test with Welch’s correction. (C) Representative images and quantification of PV^+^ puncta density in CA2/CA3 stratum pyramidale (N =4 mice per group). Top, RGS14^+^ labeling defines CA2 subfield. Scale bar, 10 μm. p < 0.05 using two-tailed unpaired *t* test with Welch’s correction (D) Images showing co-immunostaining of Dyrk1a, Ablim3 and ZO1 in hippocampus. Scale bar, 100 μm. Inset shows high magnification of dCA3 subregion outlined in image. Scale bar, 5 μm. (E) Lentiviruses expressing CaMKIIα-Cre-T2A-GFP were injected into DG of 2-month old *Dyrk1a ^f/f, f/+ and +/+^* mice 2 weeks prior to processing for morphology analysis or use in social cognition behavioral paradigm. Representative images showing GFP^+^ MFTs (filopodia extensions, arrows) and quantification of the number of vGLUT1^+^ filopodia per MFT in CA3 (left) and PV^+^ puncta density in CA3 (right)(N = 5 for mice *+/+*; 4 mice for *f/+;* 6 mice for *f/f*). Scale bar, 5 μm (left); 10 μm (right). **p < 0.01; ***p < 0.001 using one-way ANOVA with Bonferroni post hoc test. (F) Schematic of social cognition task depicting social recognition phase (Trial1, T1), two trials of familiarization or habituation to social stimulus (T2-T3) and social memory discrimination trial (T4), each trial separated by 5 min intervals. (G) Quantification of total interaction time during T1-T3. Blue shaded box outlines social recognition phase (T1). (H) Quantification of empty cup vs. social stimulus interaction time during social recognition phase (T1) (left panel, blue shaded) and the discrimination trial (T4, right panel, yellow shaded) (N =13 mice per group). Emp, empty pencil cup; stim, stimulus mouse; fam, familiar mouse; novel, a new novel stimulus mouse. **p < 0.01 using two-way ANOVA with Bonferroni post hoc test. All data are displayed as mean ± SEM. See also Figure S1.

We next asked whether *Dyrk1a* is necessary in DGCs for FFI connectivity and PV IN perisomatic contacts in CA3. *Dyrk1a* is expressed in all principal cell-types of the hippocampus (https://hipposeq.janelia.org). Immunostaining using antibodies that we validated for this study shows Dyrk1a localization in MFTs **(Figure 1D, Figure S1A)**. We injected lentiviruses expressing Cre-GFP under the control of CaMKIIα promoter into DG of adult *Dyrk1a*^+/+, f/+ or f/f^ ^f/+ or f/f^ mice (Levy et al., 2021; Thompson et al., 2015) to delete one or both alleles of *Dyrk1a* in DGCs. Loss of one or both *Dyrk1a* alleles in DGCs (referred to as *Dyrk1a^DGC f/+ or f/f^*) did not change dendritic spine densities **(Figures S1B-S1C)** but decreased the number of VGLUT1+ MFT-filopodial contacts with PV INs and PV puncta (perisomatic synaptic contacts) density in CA3 **(Figure 1E)**. Thus, normal gene dosage of *Dyrk1a* is necessary in DGCs for maintaining FFI connectivity and PV IN perisomatic contacts in CA3.

To determine if *Dyrk1a* is required in DGCs for social cognition, we tested adult wild-type mice (*Dyrk1a^DGC+/+^*) or littermates in which Dyrk1a was virally deleted just in DGCs (*Dyrk1a^DGC f/f^*) in a behavioral paradigm in which trial 1 (T1) probes social recognition, second and third trials (T2, T3) allow for familiarization and trial 4 (T4) tests discrimination of novel and familiarized social stimuli. In behavioral testing performed 2 weeks following lentiviral-Cre injections, we found that *Dyrk1a^DGC f/f^* mice did not exhibit a preference for the social stimulus over an empty cup indicating impaired social recognition. However, following familiarization trials T2 and T3, both groups of mice exhibited a comparable ability to distinguish a novel social stimulus from the familiarized social stimulus suggesting that extended exposure to social stimulus enables mice overcome deficiency in social recognition **(Figures 1F-1H)**. These data demonstrate that *Dyrka1* is necessary in DGCs of adult mice for social recognition.

Analysis of this same cohort of mice in assays for anxiety-like behavior, object preference, and novel object recognition suggested that deletion of Dyrk1a in DGCs of adult mice does not affect behavioral measures and performance in these domains (**Figures S1D-S1H**).

### Ablim3 functions downstream of Dyrk1a in mossy fibers to regulate FFI connectivity, PV IN perisomatic contacts and social recognition

We had previously shown that Ablim3 localizes to puncta adherens junctions in MFTs (visualized by immunostaining for zonula occludens 1, ZO1)-sites of stabilization of the MFT on dendritic shafts of CA3/CA2 neurons (**Figure 1D**) (Guo et al., 2018; Rollenhagen and Lubke, 2010). Since experience or viral mediated downregulation of *Ablim3* results in generation of MFT-filopodial contacts with PV INs (Guo et al., 2018; Twarkowski et al., 2022) and Dyrk1a phosphorylation of substrates may result in their degradation (De Toma et al., 2019; Schneider et al., 2015; Singh and Lauth, 2017; Thompson et al., 2015), we asked whether experience-dependent downregulation of Ablim3 in mossy fiber terminals is dependent on Dyrk1a. Towards this goal, we first generated hemizygous *Dyrk1a* mice (*EIIaCre; Dyrk1a ^f/+^* or *Dyrk1a ^+/-^*) by breeding female EIIa-Cre mice with male *Dyrk1a* ^f/+^ mice (Heffner et al., 2012; Thompson et al., 2015). EIIa-Cre mice carry a transgene under the control of the adenovirus EIIa promoter that targets expression of Cre recombinase to the preimplantation mouse embryo and are useful for germ line deletion of conditional alleles (JAX, Strain 003724) (Guo et al., 2021; Lakso et al., 1996). Analysis of EIIa-Cre activity using a Cre-reporter allele (Ai14, *B6;129S6-Gt(ROSA)26Sor^tm14(CAG-tdTomato)Hze^*/J)(Madisen et al., 2010) revealed widespread recombination in the brain (**Figure 2A**). Using *Dyrk1a* ^+/+ and +/-^ mice we quantified Ablim3 levels in MFTs following a social experience (**Figure 2A**). We found that social experience-dependent downregulation of Ablim3 was significantly reduced in MFTs of hemizygous *Dyrk1a* mice (**Figure 2B-2C**).

**Figure 2.**
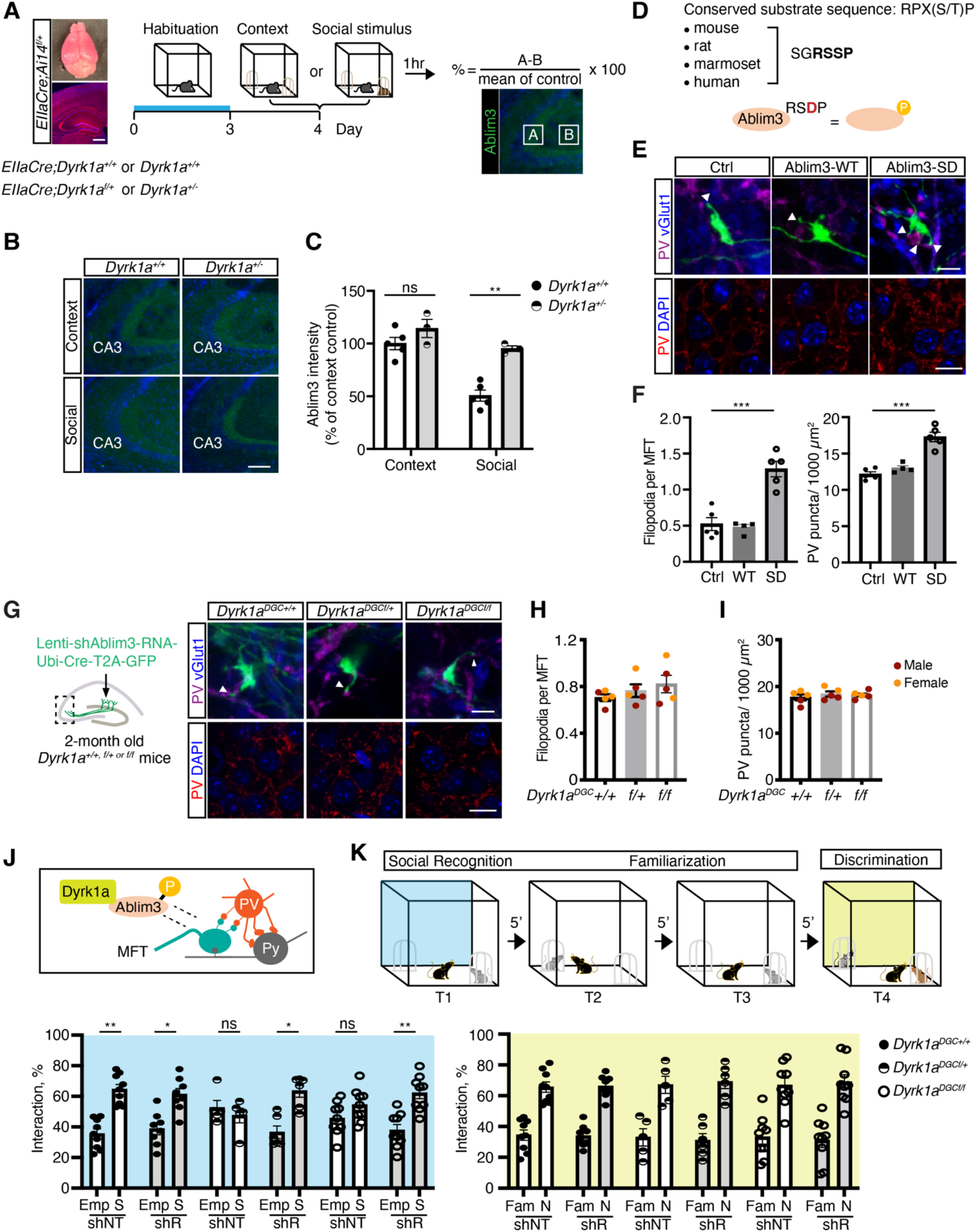
Ablim3 functions downstream of Dyrk1a in mossy fibers to regulate FFI connectivity, PV IN perisomatic contacts and social recognition. (A) Representative images of the whole brain and coronal section of *EIIaCre;Ai14* mice, schematic of experimental design and quantification of Ablim3 levels in mossy fiber terminals. Scale bar, 500 μm (B) Representative images of Ablim3 immunoreactivity in mossy fiber terminals, (referred to as *Dyrk1a^+/+ or +-^*). Scale bar, 100 μm. (C) Quantification of Ablim3 levels in mossy fiber terminals (MFTs) in 2-3 month old *EIIaCre;Dyrk1a^+/+ or f/+^* mice (referred to as *Dyrk1a^+/+ or +/-^*, in *Dyrk1a^+/+^*, N = 5 mice for each group; in *Dyrk^+/-^*, N = 3 mice for each group). **p < 0.01 using two-way ANOVA with Bonferroni post hoc test. (D) Conserved Dyrk1a phosphorylation site in Ablim3 based on consensus sequence. (E-F) Representative images of MFTs and PV puncta taken from mice following lentiviral overexpression of GFP (Ctrl), Ablim3 wild-type (WT) or phosphomimetic Ablim3 (SD) mutant in DG of 2-month old *Dyrk1a^+/+^* mice. Representative images and quantification of number of filopodia per vGLUT1^+^ MFT in CA3 (top) and PV^+^ puncta density in CA3 (bottom). Scale bar, 5 μm (top); 10 μm (bottom). (N = 5 mice for vector control; 4 mice for WT; 5 mice for SD). ***p < 0.001 using one-way ANOVA with Bonferroni post hoc test. (G) Schematic of lentivirus injection of Cre-T2A-GFP-U6-*Ablim3* shRNA cassette into DG of 2-month old *Dyrk1a^+/+ f/+ or f/f^* mice. Representative images showing GFP^+^ MFTs (filopodia extensions, arrows, top) PV^+^ puncta density in CA3 (bottom). Scale bar, 5 μm (top); 10 μm (bottom). (H-I) Quantification of the number of vGLUT1^+^ filopodia per MFT (H) and PV^+^ puncta density (I) in CA3 stratum pyramidale (N = 5 mice for each group). p > 0.05 using one-way ANOVA with Bonferroni post hoc test. (J) Model for how Dyrk1a dependent Ablim3 phosphorylation in MFTs recruits PV mediated perisomatic inhibition in CA3/CA2. (K) Schematic of social cognition task (top). Quantification of interaction time during social recognition phase (T1, left panel, blue shaded box) and social memory discrimination phase (T4, right panel, yellow shaded box). 2-month old mice injected with shNT (N = 9 mice for *+/+;* 5 mice for *f/+;* 9 mice for *f/f* mice) or shRNA (N = 8 mice for *+/+*; 6 mice for *f/+;* 9 mice for *f/f* mice). Emp, empty pencil cup; stim, stimulus mouse; fam, familiar mouse; novel, a new novel stimulus mouse. *p < 0.05; **p < 0.01 using two-way ANOVA with Bonferroni post hoc test. All data are displayed as mean ± SEM. See also Figure S2.

The Dyrk1a phosphorylation site in Ablim3 is conserved across species (**Figure 2D**). If Ablim3 functions downstream of Dyrk1a in the same pathway, then viral overexpression of an Ablim3 phosphomimetic mutant construct that mimics Dyrk1a phosphorylation of serine residue in Ablim3 in DGCs will increase FFI connectivity and PV IN perisomatic contacts in CA3 i.e. result in a gain-of-function phenotype. Viral overexpression of an Ablim3 phosphomimetic construct encoding Ablim3 with a serine to aspartic acid amino acid substitution resulted in a significant increase in VGLUT1+ MFT filopodial contacts with PV INs and PV IN perisomatic contacts in CA3 (**Figures 2E-2F**) without affecting DGC dendritic spine densities (**Figure S2A**).

Our functional epistasis analysis positions Ablim3 downstream of Dyrk1a in mossy fibers. Based on these findings, we tested whether viral downregulation of *Ablim3* in DGCs is sufficient to rescue inhibitory circuitry impairments associated with Dyrk1a LOF in DGCs in adult mice. Viral injection of a construct co-expressing Cre and GFP under the ubiquitin (*Ubi*) promoter and an extensively validated shRNA targeting Ablim3 in DGCs of adult *Dyrk1a* ^+/+, f/+ or f/f^ mice (Guo et al., 2018; Twarkowski et al., 2022) rescued Dyrk1a associated dependent reductions in VGLUT1+ MFT filopodial contacts with PV INs and PV IN puncta in CA3 (**Figures 2G-2I**) without affecting DGC dendritic spine densities (**Figure S2B**). These data support a model in which Ablim3 functions downstream of Dyrk1a in regulation of MFT-filopodial contacts with PV INs and PV IN plasticity (PV IN perisomatic contacts in CA3/CA2) (**Figure 2J)**.

We next assessed whether viral *Ablim3* downregulation in DGCs also prevents the social recognition impairment seen following loss of Dyrk1a in DGCs. Behavioral analysis of adult mice expressing non-target shRNA (shNT) or *Ablim3* shRNA (shRNA) in DGCs in which one or both *Dyrk1a* alleles were deleted showed that viral downregulation of *Ablim3* reversed the social recognition impairment (**Figure 2K**). We did not observe any effect of loss of one or both *Dyrk1a* alleles in DGCs or viral *Ablim3* downregulation in DGCs on locomotion in the open field, anxiety-like behavior, object preference or novel object recognition (**Figures S2C-2F**).

Taken together, these findings demonstrate that loss of one allele of *Dyrk1a* in DGCs results in haploinsufficiency, reduced FFI connectivity, PV IN perisomatic contacts in CA3 and impaired social recognition. Additionally, Ablim3 functions downstream of Dyrk1a in DGCs of adult mice to mediate social recognition. These acute, cell-type restricted manipulations of *Dyrk1a* and *Ablim3* motivated us to ask whether mice lacking one allele of *Dyrk1a* throughout development (hemizygous *Dyrk1a* mice) show impairments in FFI connectivity, PV IN perisomatic contacts in both CA3 and in CA2 and social recognition in adulthood.

### *Ablim3* downregulation in DGCs of adult hemizygous *Dyrk1a* mice restores FFI connectivity and PV IN perisomatic contacts in CA3 and CA2

We injected lentiviruses expressing *Ablim3* shRNA-GFP or shNT-GFP into the DG of adult *Dyrk1a* ^+/- or +/+^ mice and analyzed dendritic spine densities of DGCs, DG-PV IN-CA3/CA2 connectivity and perforant path-DGC synapses (**Figure 3A**). DGCs of adult *Dyrk1a* ^+/-^ mice (shNT group comparisons) exhibited comparable dendritic spine densities compared to DGCs of *Dyrk1a* ^+/+^ mice (**Figure S3A**) but had significantly fewer VGLUT1+ MFT filopodial contacts with PV INs and PV IN perisomatic contacts in CA3 (**Figures 3B-3C**). We noted a trend towards a significant decrease in PV IN perisomatic contacts in CA2 subregion that was labelled using the marker RGS14 (Regulator of G-protein signaling 14) (**Figure 3D**). Lentiviral downregulation of *Ablim3* in DGCs of adult *Dyrk1a* ^+/-^ mice reversed these reductions in VGLUT1+ MFT filopodial contacts with PV INs and PV IN perisomatic contacts in CA3 and in CA2 without affecting dendritic spine densities.

**Figure 3.**
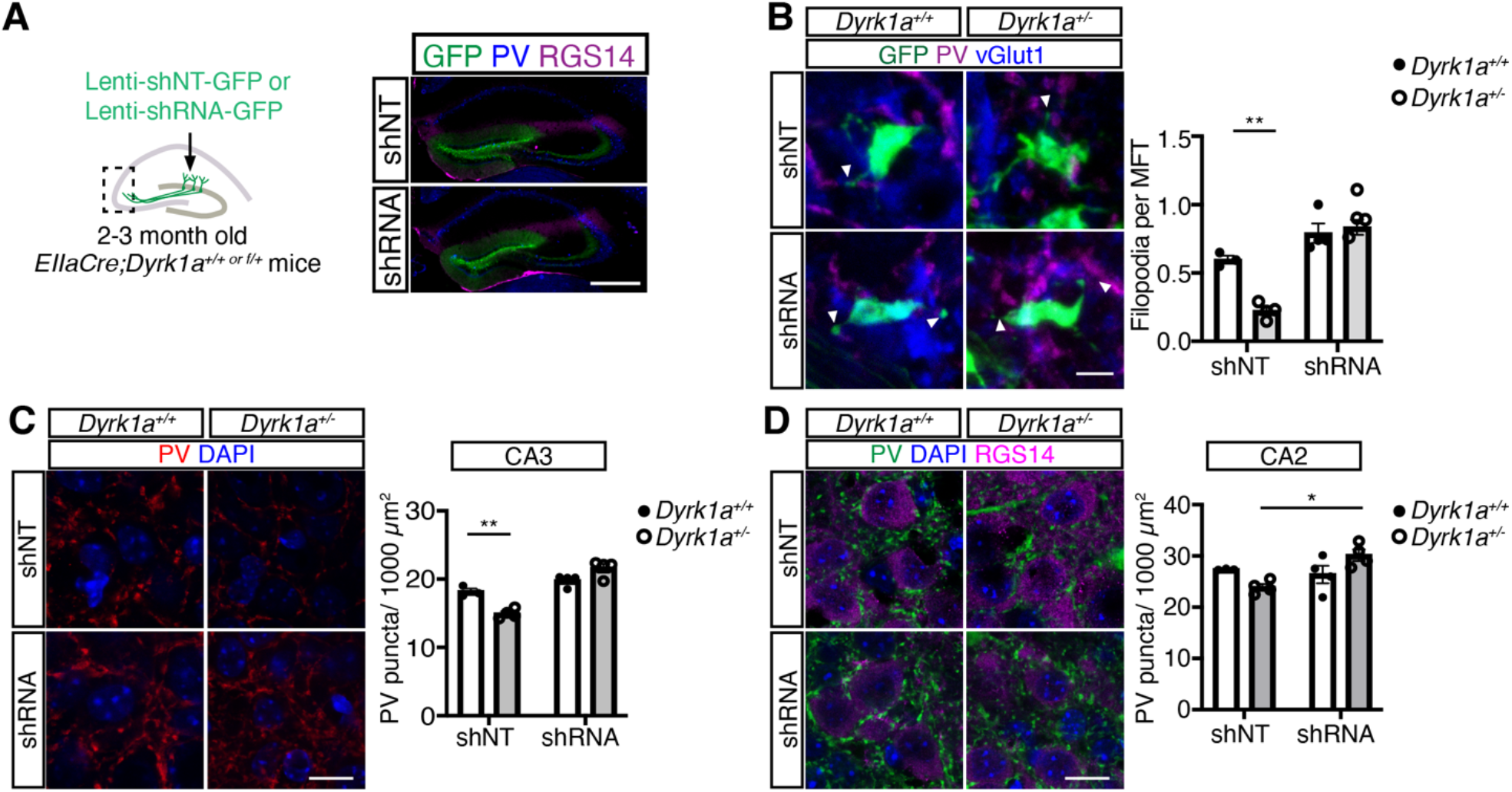
*Ablim3* downregulation in DGCs of adult hemizygous *Dyrk1a* mice restores FFI connectivity and PV IN perisomatic contacts in CA3 and CA2. (A) Lentiviruses expressing *Ablim3* shRNA-GFP or non-targeting shRNA (shNT-GFP) were injected into DG of 2-3 month old *EIIaCre;Dyrk1a^+/+ or f/+^* mice (referred to as *Dyrk1a^+/+ or +-^*). Representative images showing viral expression of GFP in DG and RGS14 immunostaining to delineate CA2. Scale bar, 500 μm. (B) Representative images showing GFP+ MFTs and quantification of the number of vGLUT1^+^ filopodia per MFT (in *Dyrk1a^+/+^*, N = 3 mice for shNT; N=4 mice for shRNA; in *Dyrk1a^f/+^*, N=4 mice for each group) in CA3. Scale bar, 5 μm. **p < 0.01 using two-way ANOVA with Bonferroni post hoc test. (C-D) Representative images and quantification of PV^+^ puncta density in CA3 (C) and CA2 (D) subfields. Scale bar, 10 μm. *p < 0.05; **p < 0.01 using two-way ANOVA with Bonferroni post hoc test. All data are displayed as mean ± SEM. See also Figure S3.

Whole-cell recordings from DGCs of adult *Dyrk1a* ^+/-^ mice did not detect a difference in frequency or amplitude of miniature excitatory postsynaptic currents (mEPSCs) (**Figure S3B**). Analysis of electrically evoked EPSCs and inhibitory postsynaptic currents (IPSCs) from DGCs revealed comparable excitation-inhibition ratios and paired-pulse ratios (PPR) for *Dyrk1a* ^+/+^ and *Dyrk1a* ^+/-^ mice (**Figure S3C**). Viral downregulation of *Ablim3* in DGCs did not affect any of these measures. These observations suggest that Dyrk1a or Ablim3 do not functionally contribute to perforant path-DGC synapses (**Figure S3C**).

Together, these findings demonstrate that viral manipulation of *Ablim3* in DGCs of adult hemizygous *Dyrk1a* mice is sufficient to reverse developmental alterations in FFI connectivity and PV perisomatic contacts in both CA3 and in CA2. We next asked whether these anatomical changes in FFI and PV IN connectivity in DG-CA3/CA2 are mirrored by electrophysiological changes in feed-forward inhibition and GABAergic inhibition of CA3 and CA2 neurons.

### *Ablim3* downregulation in DGCs of adult hemizygous *Dyrk1a* mice restores inhibitory synaptic transmission in DG-CA2

To perform *ex vivo* whole-cell recordings from PV INs in *Dyrk1a* ^+/- or +/+^ mice, we used a viral expression system, rAAV-S5E2-tdTomato, in which tdTomato expression is under the control of the “E2 regulator element” of the *Scn1a* gene (referred to as S5E2). S5E2 confers PV IN restricted expression (>90% specificity) in cortex and hippocampal CA1 of rodents (Vormstein-Schneider et al., 2020). To establish utility of using this viral expression system to selectively label PV INs in CA3/CA2, we analyzed the overlap of tdTomato and PV expression following viral injections into CA3/CA2. Systematic quantification in molecular and pyramidal layers of CA3 and CA2 revealed >90% overlap between tdTomato and PV expression (**Figure S4A**).

Next, we injected lentiviruses expressing *Ablim3* shRNA-GFP or shNT-GFP into DGCs and rAAV-S5E2-tdTomato into CA3/CA2 of adult (2-3 months) *Dyrk1a* ^+/- or +/+^ mice and performed *ex vivo* whole-cell patch-clamp recordings from CA2 PV INs and CA2 pyramidal neurons (PNs) (**Figure 4A)**. Consistent with reduced VGLUT1+ MFT filopodial contacts with PV INs in *Dyrk1a*^+/-^ mice, we observed a significant reduction in frequency, but not amplitude, of miniature excitatory postsynaptic currents (mEPSCs) in PV INs (**Figure 4B)**. In CA2 pyramidal neurons, we observed a reduction in frequency, but not amplitude, of miniature inhibitory postsynaptic currents (mIPSCs)(**Figure 4C)**. We did not detect differences in frequency or amplitude of mEPSCs in CA2 pyramidal neurons in *Dyrk1a+/-* mice (**Figure S4B**). Viral downregulation of Ablim3 in DGCs of *Dyrk1a* ^+/-^ mice completely reversed decrements in frequency of mEPSCs in PV INs in CA2 and frequency of mIPSCs in CA2 pyramidal neurons (**Figures 4B**-**4C**).

**Figure 4.**
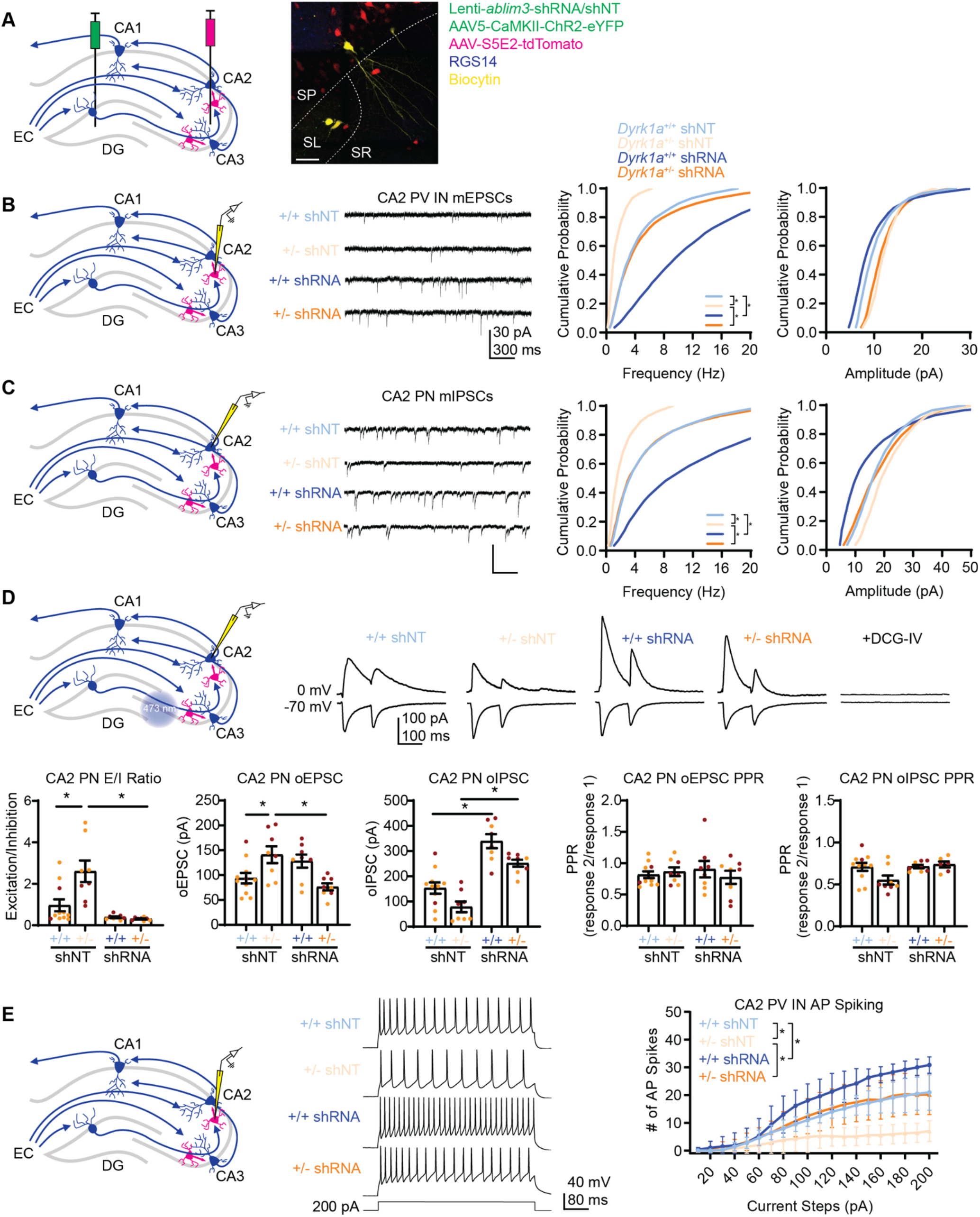
*Ablim3* downregulation in DGCs of adult hemizygous *Dyrk1a* mice restores inhibitory synaptic transmission in DG-CA2. (A) Schematic depicting lentiviral-*Ablim3*-shRNA/shNT and rAAV5-CaMKIIα-ChR2-eYFP injection into DG, and AAV-S5E2-tdTomato injected into CA3/CA2 (left). Representative image of CA2 showing eYFP expression in MFTs in stratum lucidum (SL), tdTomato expressed in PV INs, RGS14 immunostaining of CA2, and immunostaining of biocytin-filled PN and PV INs. Scale bar, 50 μm. (B) Schematic depicting whole-cell patch-clamp recording of miniature excitatory postsynaptic current (mEPSC) from CA2 PV IN (left). Representative recording traces from each group (middle). Cumulative probability plots of mEPSC frequency and amplitude from PV INs (Kolmogorov-Smirnov test, *\*p* < 0.05, n=8-9 cells, 2-3 cells per mouse, 3-4 mice per group). (C) Schematic depicting whole-cell patch-clamp recording of miniature inhibitory postsynaptic current (mIPSC) from CA2 pyramidal neurons (PN) (left). Representative traces from each group (middle). Cumulative probability plots of mIPSC frequency and amplitude from PNs (Kolmogorov-Smirnov test, \**p* < 0.05, n=8-9 cells, 2-3 cells per mouse, 3-4 mice per group). (D) Schematic depicting whole-cell patch-clamp recording of EPSC and IPSC from CA2 PN to paired pulse optical (473 nm) stimulation. Representative traces from each group and traces after perfusion of DCG-IV. Bar graphs indicate excitation to inhibition ratio, the amplitude of the first EPSC and IPSC response to paired pulse optical stimulation, and the EPSC and IPSC paired pulse ratios. (Two-way ANOVA with Tukey posthoc, \**p* < 0.05, n=8-11 cells, 2-3 cells per mouse, 3-4 mice per group). (E) Schematic depicting whole-cell patch-clamp recording of CA2 PV IN in current-clamp configuration. Representative traces from each group depict action potential response to current steps (200 pA, 500 ms). Line graph indicates number of potential responses to incremental current steps (Two-way RM ANOVA with Tukey posthoc, \**p* < 0.05, n=11-12 cells, 2-3 cells per mouse, 5-6 mice per group). See also Figure S4.

Next, we asked whether these developmental reductions in mossy fiber excitatory inputs onto PV INs and inhibitory inputs onto CA2 neurons in *Dyrk1a* ^+/-^ mice result in reduced DGC recruitment of FFI onto CA2. We virally expressed Channelrhodopsin (ChR2)(Conti, 2021) and shNT/shRNA in DGCs of *Dyrk1a* ^+/+ or +/-^ mice and recorded optically evoked excitatory and inhibitory post synaptic responses (oEPSCs and oIPSCs) in CA2 neurons following optogenetic stimulation of mossy fibers. *Dyrk1a* ^+/-^ mice exhibited a significantly higher ratio of oEPSCs and oIPSCs in CA2 pyramidal neurons compared to *Dyrk1a* ^+/+^ mice. This increase in the excitation-inhibition ratio was driven by a significant increase in oEPSCs and a reduction in oIPSCs in CA2 pyramidal neurons.Viral expression of Ablim3 shRNA in DGCs of *Dyrk1a* ^+/-^ mice resulted in complete restoration of the excitation-inhibition ratio primarily through a robust increase in oIPSCs in CA2 pyramidal neurons. Bath application of DCG-IV, a group II metabotropic glutamate receptor agonist that selectively blocks evoked MFT release, to hippocampal slices abolished optically evoked inhibitory and excitatory responses in CA2 pyramidal neurons. We did not detect changes in oEPSC or oIPSC paired pulse ratios (PPRs) indicating that the vesicle release machinery in MFTs was not affected in *Dyrk1a* ^+/-^ mice or following *Ablim3* downregulation in DGCs (**Figure 4D**).

We next assessed changes in intrinsic excitability of PV INs in CA2 in *Dyrk1a* ^+/-^ mice. Current injections in PV INs revealed reduced excitability in *Dyrk1a* ^+/-^ mice which was rescued by *Ablim3* downregulation in DGCs (**Figure 4E**). We did not detect significant differences in passive membrane properties of PV INs between *Dyrk1a* genotypes. However, *Ablim3* downregulation in DGCs reduced the threshold for action potential in PV INs (**Figure S4C**). These findings suggest that increasing mossy fiber excitatory drive onto PV INs restored intrinsic excitability of these cells.

Taken together, these findings demonstrate that viral *Ablim3* downregulation in DGCs in adulthood reverses *Dyrk1a* hemizygosity-associated developmental impairments in FFI in DG-CA2, CA2 PV IN intrinsic excitability, and GABAergic inhibition in CA2. We next investigated inhibitory synaptic transmission in DG-PV IN-CA3 circuitry in *Dyrk1a* ^+/-^ mice.

### *Ablim3* downregulation in DGCs of adult hemizygous *Dyrk1a* mice restores inhibitory synaptic transmission in DG-CA3

Whole-cell recordings from tdTomato labeled PV INs in CA3 of *Dyrk1a^+/-^* mice revealed a significant reduction in frequency, but not amplitude, of mEPSCs (**Figures 5A-5B)**. Consistent with reduced PV IN perisomatic contacts in CA3, we observed a reduction in frequency, but not amplitude, of mIPSCs in CA3 pyramidal neurons (**Figure 5C)**. We did not detect differences in frequency or amplitude of mEPSCs in CA3 pyramidal neurons in *Dyrk1a+/-* mice (**Figure S5A**). Viral downregulation of Ablim3 in DGCs of *Dyrk1a* ^+/-^ mice completely reversed decrements in frequency of mEPSCs in PV INs in CA3 and frequency of mIPSCs in CA3 pyramidal neurons (**Figures 5B**-**5C**).

**Figure 5.**
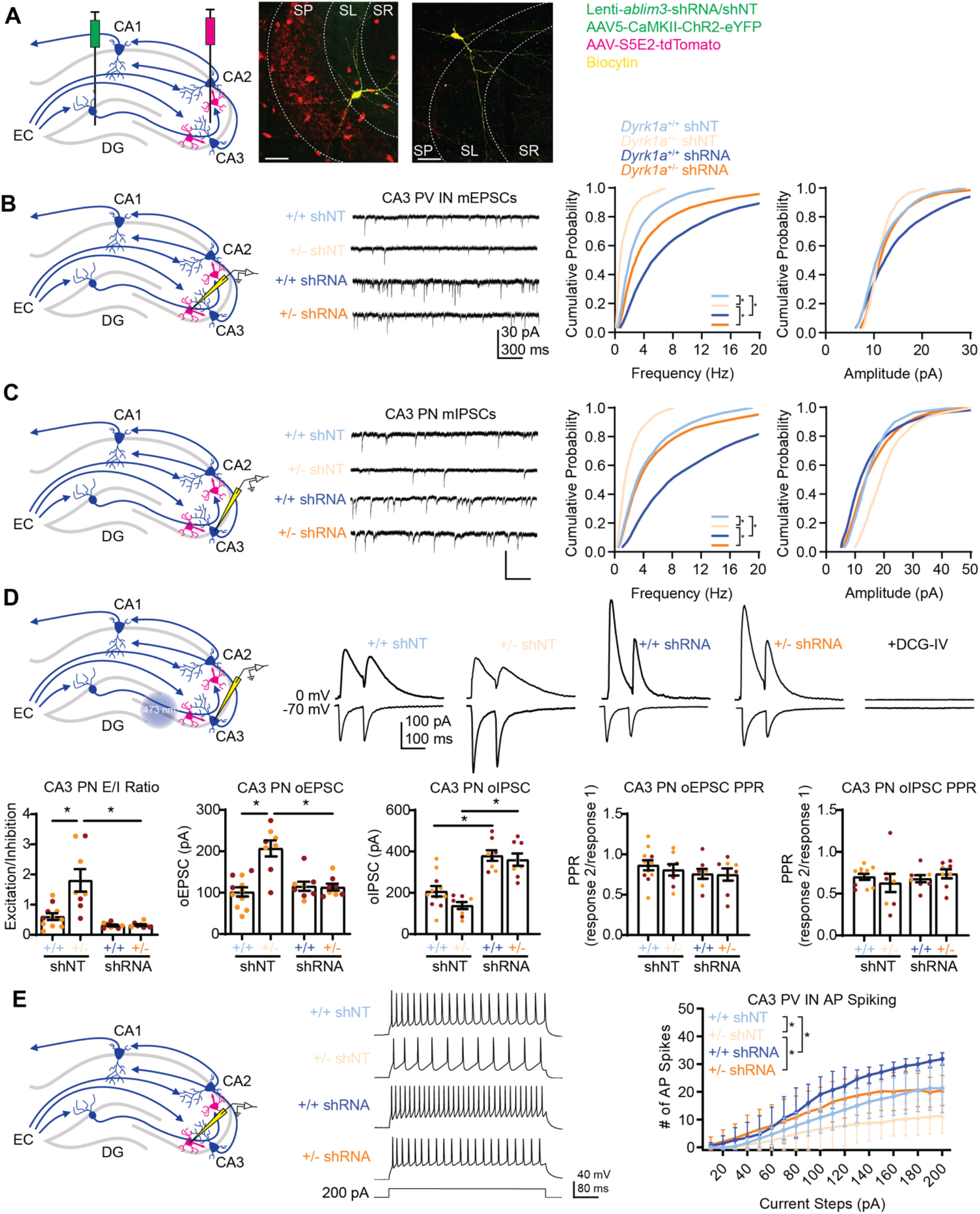
*Ablim3* downregulation in DGCs of adult hemizygous *Dyrk1a* mice restores inhibitory synaptic transmission in DG-CA3. (A) Schematic depicting lentiviral-ablim3-shRNA/shNT and rAAV5-CaMKIIα-ChR2-eYFP injection into DG, and AAV-S5E2-tdTomato injected into CA3/CA2 (left). Representative image of CA2 showing eYFP expression in MFTs in the stratum lucidum (SL), tdTomato expressed in PV^+^ INs, and immunostaining of biocytin-filled PN and PV^+^ INs. Scale bar, 50 μm. (B) Schematic depicting whole-cell patch-clamp recording of mEPSC from CA3 PV IN (left). Representative recording traces from each group (middle). Cumulative probability plots of mEPSC frequency and amplitude from PV INs (Kolmogorov-Smirnov test, *\*p* < 0.05, n=8-9 cells, 2-3 cells per mouse, 3-4 mice per group). (C) Schematic depicting whole-cell patch-clamp recording of mIPSC from CA3 PN (left). Representative traces from each group (middle). Cumulative probability plots of mIPSC frequency and amplitude from PNs (Kolmogorov-Smirnov test, \**p* < 0.05, n=8-9 cells, 2-3 cells per mouse, 3-4 mice per group). (D) Schematic depicting whole-cell patch-clamp recording of EPSC and IPSC from CA3 PN to paired pulse optical (473 nm) stimulation. Representative traces from each group and traces after perfusion of DCG-IV. Bar graphs indicate excitation to inhibition ratio, the amplitude of the first EPSC and IPSC response to paired pulse optical stimulation, and the EPSC and IPSC paired pulse ratios. (Two-way ANOVA with Tukey posthoc, \**p* < 0.05, n=8-11 cells, 2-3 cells per mouse, 3-4 mice per group). (E) Schematic depicting whole-cell patch-clamp recording of CA3 PV^+^ IN in current-clamp configuration. Representative traces from each group depict action potential response to current steps (200 pA, 500 ms). Line graph indicates number of potential responses to incremental current steps (Two-way RM ANOVA with Tukey posthoc, \**p* < 0.05, n=13-13 cells, 2-3 cells per mouse, 5-7 mice per group). See also Figure S5.

Next, we asked whether these developmental reductions in excitatory inputs onto PV INs and inhibitory inputs onto CA3 neurons in *Dyrk1a* ^+/-^ mice result in reduced DGC recruitment of FFI onto CA3. Analysis of oEPSCs and oIPSCs in CA3 neurons following optogenetic stimulation of mossy fibers revealed a significantly higher excitation-inhibition ratio in *Dyrk1a* ^+/-^ mice that was driven by a significant increase in oEPSCs (**Figure 5D**). Viral expression of Ablim3 shRNA in DGCs of *Dyrk1a* ^+/-^ mice resulted in complete restoration of the excitation-inhibition ratio primarily through a robust increase in oIPSCs in CA3 pyramidal neurons. Bath application of DCG-IV to hippocampal slices abolished optically evoked inhibitory and excitatory responses in CA3 pyramidal neurons. We did not detect any changes in oEPSC and oIPSC PPRs in CA3 neurons of *Dyrk1a* ^+/-^ mice or following *Ablim3* downregulation in DGCs (**Figure 5D**).

As with PV INs in CA2, current injections in PV INs in CA3 revealed reduced intrinsic excitability in *Dyrk1a* ^+/-^ mice which was rescued by *Ablim3* downregulation in DGCs (**Figure 5E**). We did not detect significant differences in passive membrane properties of PV INs between *Dyrk1a* genotypes (**Figure S5B**).

Taken together, these findings demonstrate that viral *Ablim3* downregulation in DGCs in adulthood reverses *Dyrk1a* hemizygosity-associated developmental impairments in FFI in DG-CA3, CA3 PV IN intrinsic excitability and GABAergic inhibition in CA3.

### *Ablim3* downregulation in DGCs of adult hemizygous *Dyrk1a* mice restores social recognition

To determine if *Dyrk1a* hemizygosity impairs social cognition and whether *Ablim3* downregulation in DGCs may exert restorative effects, we tested adult wild-type mice or *Dyrk1a^+/-^* littermates expressing shRNA or shNT in DGCs in the social recognition and discrimination task (**Figure 6A**). *Dyrk1a ^+/-^* mice were impaired in social recognition i.e they spent equivalent amount of time investigating the social stimulus and empty cup in the first trial but learned to distinguish a novel social stimulus from the familiarized social stimulus in trial 4 **(Figure 6F)**. Viral downregulation of *Ablim3* in DGCs of *Dyrk1a ^+/-^* mice rescued this impairment in social recognition. Analysis of this same cohort of mice in assays for anxiety-like behavior, object preference, and novel object recognition suggested that *Dyrk1a* hemizygosity does not affect behavioral measures and performance in these domains (**Figures 6B-6E**). Interestingly, developmental deletion of *Dyrk1a* in hippocampus and cortex also produced a similar impairment in social recognition without affecting anxiety-like behavior (Levy et al., 2021). Taken together, our findings demonstrate that viral *Ablim3* downregulation in DGCs in adulthood is sufficient to reverse *Dyrk1a* hemizygosity-associated developmental impairments in social recognition.

**Figure 6.**
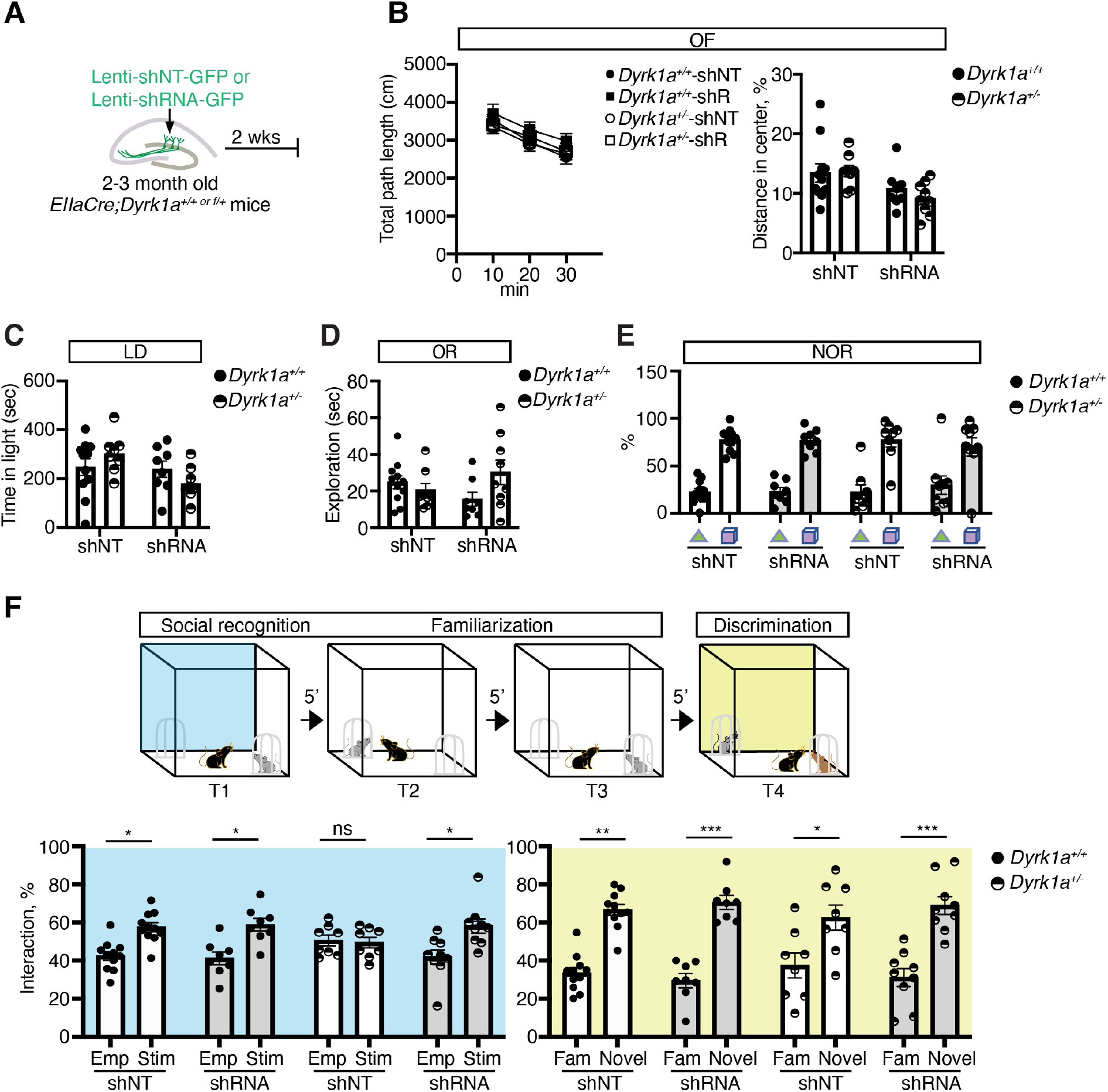
*Ablim3* downregulation in DGCs of adult hemizygous *Dyrk1a* mice restores social recognition. (A) Lentiviruses expressing *Ablim3* shRNA-GFP or non-targeting shRNA (shNT-GFP) were injected into DG of 2-3 month old *EIIaCre;Dyrk1a^+/+ or f/+^* mice (referred to as *Dyrk1a^+/+ or +-^*). (B-F) Behavioral analysis of *Dyrk1a^+/+^* mice injected with shNT (N=11 mice) or shRNA (N=8 mice) and *Dyrk1a ^+/-^* mice injected with shNT (N=8 mice) or shRNA (N=9 mice)(see Figure S1D for behavioral testing schedule). (B) OF, open-field, quantification of total distance travelled (left) and percentage of distance travelled across the center arena (right). (C) LD, light-dark box assay, quantification of time spent in the light compartment (seconds). (D) OR, object recognition, quantification of the time spent sniffing the object (seconds). (E) NOR, novel object recognition, quantification of the time spent sniffing the familiar object (triangle) and novel object (cube). (F) Quantification of cup vs. social stimulus interaction time during social recognition phase (T1) (left panel, blue shaded) and the discrimination trial (T4, right panel, yellow shaded) *p < 0.05; **p < 0.01; ***p < 0.001 using two-way ANOVA with Bonferroni post hoc test. All data are displayed as mean ± SEM.

### Acute chemogenetic activation of PV INs in CA3/CA2 of adult *Dyrk1a*^+/-^ mice is sufficient to rescue social recognition

So far, our data identify Ablim3 and PV IN mediated perisomatic inhibition as molecular and circuit substrates of Dyrk1a function in DG-CA3 and DG-CA2 circuits. We next investigated the potential of directly activating PV INs in CA3/CA2 in *Dyrk1a* ^+/-^ mice to rescue the social recognition impairment. We performed stereotaxic injections of rAAV-S5E2-hM3D(Gq)DREADD-P2A-dTomato into CA3/CA2 of *Dyrk1a*^+/+ or +/-^ mice to express the chemogenetic activator hM3D(Gq)-DREADD (Roth, 2016) exclusively in PV INs (**Figure 7A**). Using a cross-over CNO/saline design with adult male and female *Dyrk1a* ^+/+ or +/-^ mice, we found that acute chemogenetic activation of PV INs in CA3/CA2 did not affect anxiety-like behavior, object preference, and novel object recognition (**Figures 7B-7D**) but was sufficient to rescue social recognition impairments (**Figure 7E**). Acute CNO treatment as assessed using *Dyrk1a* ^+/+^ mice did not affect anxiety-like behavior, object recognition or social recognition behavior (**Figures S6A-6E**).

**Figure 7.**
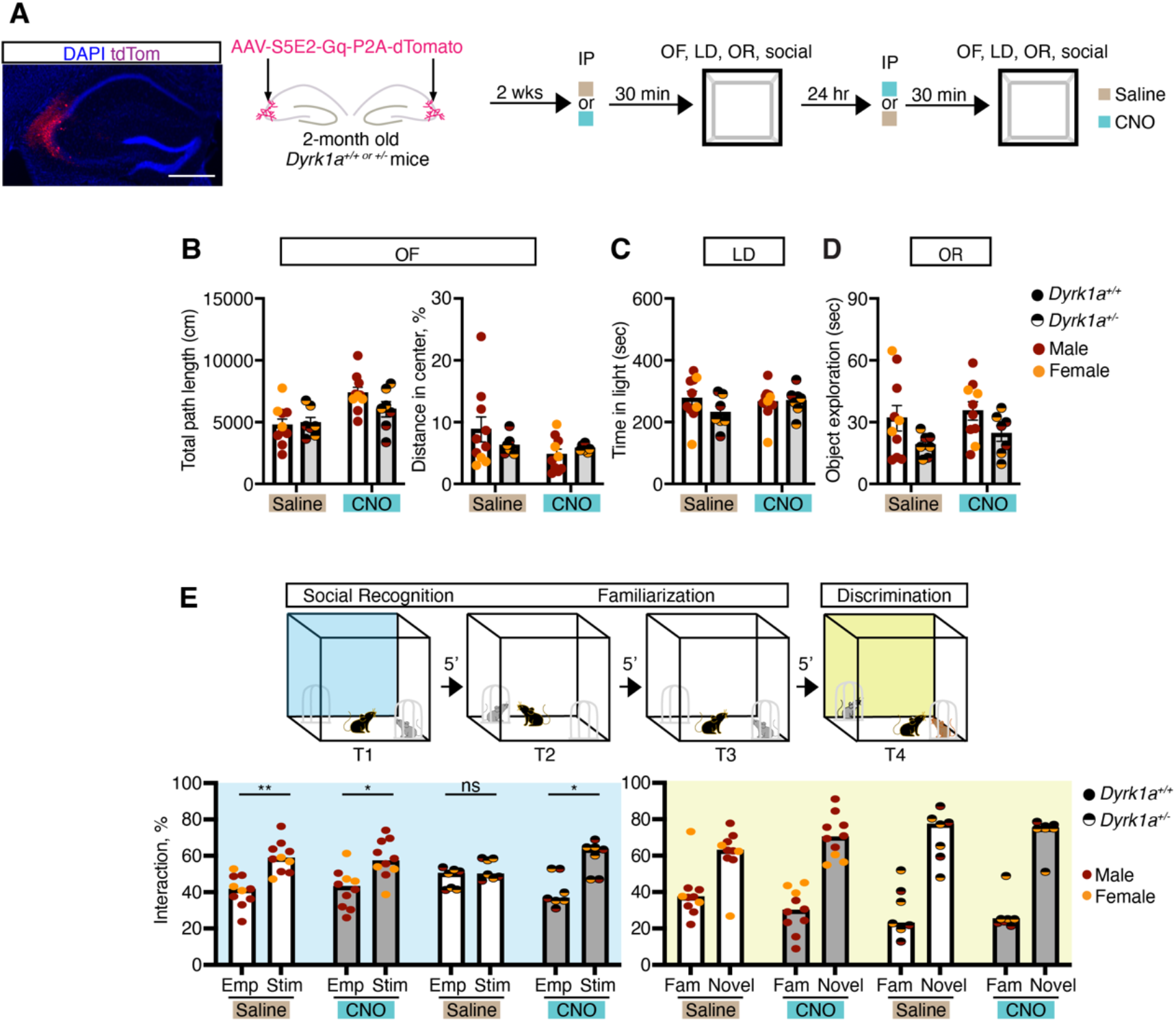
Acute chemogenetic activation of PV INs in CA3/CA2 of adult *Dyrk1a^+/-^* mice is sufficient to rescue social recognition. (A) Left, Representative image showing expression of hM3D(Gq)DREADD-tdTom in PV INs in CA3/CA2. Scale bar, 500 μm. Right, Schematic of chemogenetic activation of PV INs schedule. 2-month old *EIIaCre;Dyrk1a^+/+ or f/+^* mice were injected with AAV-S5E2-Gq virus into dCA2/CA3 2 weeks prior to behavioral testing using a vehicle/CNO cross-over design. Mice were injected with saline or 1mg/kg CNO 30 min prior to behavioral testing. 24 hours later mice were counterbalanced for vehicle/CNO and behaviorally tested. (B-D) Chemogenetic activation of PV INs in CA3/CA2 of does not affect locomotion (B), anxiety-like behavior (C) and object recognition (D). (E) Quantification of cup vs. social stimulus interaction time during social recognition phase (T1) (left panel, blue shaded) and the discrimination trial (T4, right panel, yellow shaded) (N= 10 mice for *Dyrk1a^+/+^* group; N= 7 mice for *Dyrk1a^+/-^* group). *p < 0.05; **p < 0.01; ***p < 0.001 using two-way ANOVA with Bonferroni post hoc test. All data are displayed as mean ± SEM. See also Figure S6.

## DISCUSSION

*Dyrk1a* haploinsufficiency is associated with a constellation of somatic, neurological and neurobehavioral symptoms including ASD. A major challenge is understanding how *Dyrk1a* hemizygosity impairs social cognition. Studies in invertebrates and mice have found that *Dyrk1a* LOF mutations affect dendritic arborization, soma size, axonal elongation, dendritic spines, presynaptic terminal growth and synaptic vesicle recycling (Arranz et al., 2019; Atas-Ozcan et al., 2021; Benavides-Piccione et al., 2005; Chen et al., 2014; Classen et al., 2020; Dang et al., 2018; Duchon and Herault, 2016; Fotaki et al., 2002; Levy et al., 2021; Soppa and Becker, 2015; Souchet et al., 2014; Thompson et al., 2015; Vidaki et al., 2017). Additionally, frameshift mutations in *Dyrk1a* (Raveau et al., 2018) or conditional deletion of *Dyrk1a* in neocortex and hippocampus (Levy et al., 2021) impair social recognition in mice. However, despite this knowledge, we do not know how Dyrk1a regulates synaptic and circuit functions to influence social cognition. Here, we demonstrate a role for Dyrk1a in DGCs in regulating mossy fiber recruitment of PV INs, maintaining PV IN intrinsic excitability and recruiting GABAergic inhibition of CA3 and CA2. We show that acute loss of one functional allele of *Dyrk1a* in DGCs is sufficient to reduce MFT-filopodial contacts with PV INs and PV IN perisomatic contacts in CA3. Global loss of a functional allele of *Dyrk1a* throughout life also resulted in reduced mossy fiber recruitment of PV INs and decreased GABAergic inhibition of CA3 and CA2 without affecting DGC dendritic spine density, PP-DGC synaptic properties or mossy fiber vesicle release probability. Acute loss of a functional allele of *Dyrk1a* in DGCs or global *Dyrk1a* hemizygosity resulted in impaired social recognition. Together, these acute, cell-type specific and global, developmental manipulations of *Dyrk1a* identify a dosage sensitive function for Dyrk1a in DGCs in recruiting PV IN mediated GABAergic inhibition in CA3 and CA2 and regulation of social recognition.

In the absence of known activators of Dyrk1a function, can we devise strategies to reverse *Dyrk1a* haploinsufficiency-associated reductions in DGC recruitment of FFI of CA3/CA2 and social recognition impairment? Here, we implemented functional epistasis logic to target a downstream effector and synaptic substrate of Dyrk1a in mossy fibers, Ablim3, to achieve this goal. In prior work, we showed that Ablim3 functions as a molecular brake of DGC mediated FFI in DG-CA3. Here, we generated multiple lines of evidence to demonstrate that Ablim3 functions downstream of Dyrk1a in mossy fibers to recruit FFI in DG-CA2 and DG-CA3. First, we found that hemizygous *Dyrk1a* mice failed to show social experiencedependent Ablim3 downregulation in mossy fibers. Second, viral expression of an Ablim3 phosphomimetic mutant (that mimics Dyrk1a phosphorylation of Ablim3) in DGCs of wild-type mice resulted in increased MFT-filopodial contacts with PV INs and PV IN perisomatic contacts with CA3/CA2 neurons. Third, we found that viral downregulation of Ablim3 in DGCs lacking one or both alleles of *Dyrk1a* was sufficient to restore FFI connectivity, PV IN perisomatic contacts and social recognition. Building on these findings, we showed that viral Ablim3 downregulation in DGCs of adult hemizygous *Dyrk1a* mice was sufficient to restore DGC recruitment of GABAergic inhibition in CA3 and CA2, intrinsic excitability of PV INs and social recognition.

Our study implicates neural activity and more specifically, mossy fiber excitatory inputs, in regulating PV IN mediated perisomatic innervation and GABAergic inhibition of CA3 and CA2 neurons. These observations suggest that PV INs in the adult brain continue to exhibit experience-dependent plasticity of perisomatic inhibition that is observed during sensitive periods of brain development (Chattopadhyaya et al., 2004; McFarlan et al., 2023). Interestingly, we also found that increasing mossy fiber excitatory drive onto PV INs increased PV IN intrinsic excitability or intrinsic plasticity (McFarlan et al., 2023). Thus, mossy fiber recruitment of FFI in DG-CA3/CA2 circuits may also depend on increased PV IN excitability as shown for FFI in CA3-CA1 circuits (Campanac et al., 2013). Whether developmental, genetic or epigenetic programs involved in PV IN maturation, excitability and specification of synaptic connectivity are re-used to mediate activity-dependent changes in PV INs excitability and perisomatic inhibition in DG-CA3/CA2 circuits is to be determined (Allaway et al., 2021; Bernard et al., 2022; Dehorter et al., 2015; Exposito-Alonso and Rico, 2022; Favuzzi et al., 2019; Li et al., 2011; Uezu et al., 2016).

A growing body of work implicates entorhinal cortex-DG-CA3/CA2 circuits in different facets of social cognition such as social recognition, social memory discrimination and consolidation (Alexander et al., 2018; Alexander et al., 2016; Chen et al., 2020; Chiang et al., 2018; Cope et al., 2020; Deng et al., 2019; Finlay et al., 2015; Hitti and Siegelbaum, 2014; Lehr et al., 2021; Leung et al., 2018; Li et al., 2022; Lin et al., 2018; Meira et al., 2018; Oliva, 2022; Oliva et al., 2016; Oliva et al., 2020; Raam et al., 2017; Stevenson and Caldwell, 2014). This study causally links a hippocampal circuit mechanism, DGC recruitment of PV mediated inhibition of CA3 and CA2, with social recognition. We show that mossy fiber excitatory inputs onto PV INs and PV IN-CA3/CA2 inhibitory circuits are highly sensitive to social experience. Restoration of PV IN mediated inhibition onto CA3 and CA2 by either increasing excitatory drive onto PV INs and enhancing PV IN plasticity (inhibitory contacts with CA3/CA2 neurons) or direct chemogenetic activation of PV INs is sufficient to restore social recognition in adult hemizygous *Dyrk1a* mice. How does PV mediated perisomatic inhibition of CA3/CA2 contribute to social recognition? Perisomatic inhibition of CA3/CA2 neurons may impose a sparse activity regimen that supports generation of distinct neuronal ensembles in response to an experience (Csicsvari et al., 2000; de la Prida et al., 2006; Gómez-Ocádiz et al., 2021; Guo et al., 2018; Mori et al., 2007; Neubrandt et al., 2017; Sasaki et al., 2018; Torborg et al., 2010; Twarkowski et al., 2022). Consistent with this idea, DGC recruitment of FFI in CA3/CA2 is randomly wired so as to provide blanket inhibition and govern network excitability in CA3 and CA2, rather than couple individual DGC-dependent excitation with inhibition onto distinct populations of pyramidal neurons (Neubrandt et al., 2017). Loss of PV IN mediated inhibition may disrupt neuronal ensembles and network oscillations by impairing neuronal spiking, recurrent excitation in CA3 networks (Sadeh and Clopath, 2021), reciprocal inhibition between CA3 and CA2 (Boehringer et al., 2017; Fernandez-Lamo et al., 2019; Lehr et al., 2021; Middleton and McHugh, 2020; Nasrallah et al., 2019; Stober et al., 2020) and/or the balance between subcortical and entorhinal inputs to CA3/CA2 during encoding of social stimuli (Chen et al., 2020; Lopez-Rojas et al., 2022; Robert et al., 2021; Wu et al., 2021). Future studies will edify how PV inhibition of CA3/CA2 facilitates encoding of social stimuli in CA3 and CA2 neuronal ensembles and network oscillations.

Our work exemplifies how functional epistasis logic can be deployed to rescue *Dyrk1a* haploinsufficiency associated circuit and behavioral impairments. Such an approach is advantageous over efforts to identify Dyrk1a activators for several reasons. First, screening and identification of activators of kinases is notoriously difficult and currently, there are no known chemical activators of Dyrk1a. Thus, targeting specific Dyrk1a substrates in distinct somatic, neuronal and non-neuronal cell-types that mediate different Dyrk1a functions in physiology and behavior may represent a more viable approach. Second, *Ablim3*, like other Dyrk1a substrates shows more cell-type and tissue restricted expression than *Dyrk1a*. Consequently, targeting Dyrk1a substrates may permit rescue of subsets of somatic and neurological endophenotypes that make up syndromic ASD while reducing risk of cancer malignancies and neuropathology associated with widespread Dyrk1a overactivation (Lee et al., 2021; Ruiz-Mejias et al., 2016; Souchet et al., 2014; Thompson et al., 2015).

While it remains challenging to precisely determine how *Dyrk1a* haploinsufficiency affects the trajectory of brain development or for that matter, restore the arc of “normal” development, our work illuminates the potential for targeting Dyrk1a synaptic substrates in adulthood to alleviate the burden of developmentally accrued impairments in circuitry and cognition associated with *Dyrk1a* hemizygosity. Indeed, a small but growing number of studies support the promise of restoring functions in neurodevelopmental disorders after sensitive periods of brain development (Ehninger et al., 2008; Guy et al., 2007; Mielnik et al., 2021; Sztainberg et al., 2015). The last few years has witnessed the emergence of gene therapy as a potential therapeutic modality for neurological and psychiatric diseases (Davidson et al., 2022). This paradigm shift in thinking and approach to brain diseases has been catalyzed by advances in directed capsid evolution and AAV engineering, cell-type specific payload expression such as that shown here for PV INs (Vormstein-Schneider et al., 2020), antisense oligo modifications (intrathecal delivery and intravenous) and gene delivery modalities (Dunbar et al., 2018; Goertsen et al., 2021; Nagata et al., 2021; Roberts et al., 2020; Segel et al., 2021; Sun and Roy, 2021). Targeting molecular effectors of Dyrk1a such as Ablim3 using antisense oligo, siRNA or viral shRNA gene therapy or viral mediated enhancement of PV IN mediated inhibition of CA3/CA2 may harbor potential for ameliorating social recognition impairments associated with *Dyrk1a* haploinsufficiency and potentially, other neurodevelopmental disorders characterized by deficits in FFI in DG-CA3/CA2 circuits and PV IN dysfunction (Exposito-Alonso and Rico, 2022; Martin et al., 2015; Willsey et al., 2022).

## METHODS

### Subjects

All mice were group housed and experiments were conducted in accordance with procedures approved by the Institutional Animal Care and Use Committees at the Massachusetts General Hospital and NIH guidelines (IACUC 2011N000084). All mice were housed in a 12-h (7 a.m. to 7 p.m.) light–dark colony room at 22–24 °C with *ad libitum* access to food and water. Adult female mice (3–4 months old) were purchased from the Jackson Laboratories for breeding. Mouse lines were obtained from the Jackson Laboratories and details are provided in the resource table.

### Viruses and virus constructs

Lenti-shNT and shRNA (*Ablim3*) viruses were generated in the laboratory as described previously (Guo et al., 2018). rAAV viruses were purchased from Addgene. Lenti-shNT-Ubi-Cre-T2A-GFP and shRNA-Ubi-Cre-T2A-GFP (*Ablim3*) constructs were generated by subcloning Cre-T2A-GFP from plasmid-CamKIIα-Cre-T2A-GFP (VectorBuilder) into lenti-shNT and shRNA (*Ablim3*) plasmids. Lentiviruses were produced by cotransfection of lenti vectors and packaging plasmids into HEK293T cells. Briefly, 24 hours following transfection, the supernatant was collected every day for 3 days. The supernatant was concentrated by ultra-centrifugation. Virus pellets were resuspended overnight in Dulbecco’s phosphate-buffered saline (DPBS) (Guo et al., 2018).

### Stereotactic viral injection

Mice received carprofen (5mg/kg subcutaneously, Patterson Veterinary Supply) before surgery and were then anaesthetized with ketamine and xylazine (10 mg/ml and 1.6 mg/ml, IP). Mice were placed in the stereotaxic apparatus, and a small hole was drilled at each injection location (Foredom K.1070 High Speed Rotary Micromotor Kit). Bilateral injections were performed using Hamilton microsyringes (Hamilton, Neuros Syringe 7001) that were slowly lowered into target sites and that were left there for 8 min prior to infusion at a rate of 0.1 μl/min. The coordinates relative to bregma: dorsal DG: −1.8 mm (AP), ±1.35 mm (ML), −2.25 mm (DV) and dorsal CA2/CA3: −1.8 mm (AP), ±2.45 mm (ML), −2.35 mm (DV). The microsyringes were slowly withdrawn after infusion and the skin above the incision was sutured with coated vicryl sutures (Ethicon US LLC). Mice were monitored and received a daily injection of carprofen (5 mg/kg, i.p) for 3 days following surgery (Twarkowski et al., 2022).

### Behavioral procedures

Two weeks after viral injections, mice were handled for 3 days prior to behavioral experiments to habituate them to human handling and transportation from vivarium to behavioral testing rooms. For chemogenetic actuation of PV INs, mice were habituated to an empty syringe for 3 days and then CNO (1mg/kg) was injected 30 min prior to experiments. The behavioral assays were performed in the following order: open-field (OF, day 1), light-dark test (day 2), object recognition (OR, day 3), novel object recognition (NOR, day 4); social recognition and discrimination (day 6). For assessing effects of context vs. social experience on FFI connectivity, mice were habituated to the behavioral chamber for 7 days. All the behavioral assays were performed in the same chambers (40 × 40 cm, MazeEngineers). Videos were recorded and exported from Freezeframe (Actimetrics) and analyzed with EthoVision XT 15 (Noldus). Center point tracking was used to record movement and nose point tracking was used to evaluate object and social interaction. An interaction was considered when the test mouse’s nose position to object or stimulus mouse was within 1 cm. (Guo et al., 2018; Twarkowski et al., 2022).

### Open-field paradigm

Mice were transported into a holding room and habituated for one hour prior to testing. Total distance traveled and the time spent in the center of the arena were quantified over 30 min (Guo et al., 2018)..

### Light-Dark task

A dark opaque box (40 x 20 cm) with an opening in the center of one wall was placed in the OF chamber to create light and dark compartments of equal size. The mouse was placed in the light compartment and allowed to explore freely for 10 min. Time spent in the light compartment was recorded (Guo et al., 2018; Raam et al., 2017).

### Object recognition

One object (4 x 4 x 6 cm) was placed in the OF chamber along one side (5 cm distance from the wall). Mice were placed in the opposite side of the object and allowed to explore freely for 5 min. Time spent exploring the object (noise point within 2 cm) was quantified (Guo et al., 2018).

### Novel object recognition

One object (2 x 4 x 6 cm) was placed in the OF chamber along one corner and the other one was placed in the opposite corner (2 x 2 x 10 cm, 5 cm distance from the wall). The mouse was placed in the center and allowed to explore freely for 5 min. Time spent exploring the objects (nose point within 2 cm) were quantified (Guo et al., 2018; Raam et al., 2017).

### Social recognition, familiarization and social discrimination

Stimulus mice (strain, age and sex-matched) were habituated to being placed in a pencil wire cup in the OF chamber prior to the task day for 3 days, 15 min per day. The task consisted of 3 trials for familiarization and one trial for discrimination with 5 min intertrial intervals. For trial 1 (social recognition), subjects were placed in the center of the chamber for 10 min with two pencil wire cups placed on opposite corners. A beaker was placed above the pencil cup to prevent subject from climbing on the top. One cup contained a stimulus mouse; the other one remained empty. Subjects were trained from trial 1 (social recognition) to trial 3 to familiarize the stimulus mouse with the test mouse. The locations of the stimulus and empty cup were counterbalanced across trials. On trial 4 (discrimination), a novel stimulus mouse was added to the empty cup. Time spent exploring the empty and stimulus mouse were quantified (nose point within 1 cm)(Raam et al., 2017).

### Immunohistochemistry

Mice were anaesthetized with ketamine and xylazine (10 mg/ml and 1.6 mg/ml, IP), transcardially perfused with 4% PFA, and brains were incubated in 4% PFA at 4°C overnight. Brains were placed in 30% sucrose/PBS for 2 days and then embedded in medium (OCT, Fisher HealthCare). 35 μm cryosections were obtained (Leica) and stored in PBS (0.01% sodium azide) at 4°C. For immunostaining, floating sections were permeabilized, blocked in blocking solution for 2 h (PBS containing 0.3 % Triton X-100 and 10% normal donkey serum, NDS), and followed by incubation with primary antibodies (PBS containing 10% NDS) at 4°C overnight. Sections were then washed with PBS 3 times, 10 min each, followed by incubated with secondary antibodies in PBS for 2 h at room temperature (RT). Sections were then washed with PBS 3 times, 10 min each, mounted on glass slides and cover slipped with mounting medium containing DAPI. *For Dyrk1a and Ablim3 immunostaining*, brains were rapidly removed and flash frozen. 20 μm cryosections were mounted on glass slides, air-dried, and fixed in 95% ethanol immediately at −20°C for 30 min, followed by 100% acetone for 1 min at RT. Slides were rinsed with PBS, blocked with 1% BSA for 30 min, then incubated with primary antibodies (in blocking buffer) for 2 h. Slides were washed with PBS 3 times, 5 min each, sections were incubated with secondary antibodies (in PBS) for 30 min. Finally, sections were washed with PBS 3 times, 5 min each, and cover slipped with mounting medium containing DAPI. See Key Resource Table for primary antibodies information and dilutions.

### Image analysis of mossy fiber terminals, PV puncta and dendritic spines

Images were obtained from 6 sections per mouse hippocampus. A Leica SP8 confocal laser microscope and LAS software were used to capture images in the stratum lucidum at high-resolution (2,048). For MFTs, images were captured in the CA3ab subfield with a 0.3 μm step size of *z*-stacks using a 63× oil objective plus 6× digital zoom. MFT filopodia were defined as protrusion longer than 1 μm with an end-swelling structure; for PV puncta, single confocal plane image was captured in the CA2 and CA3ab subfields using a 63× oil objective plus 4× digital zoom; for dendritic spines, images were captured in the outer one-third of the molecular layer of the DG with a 0.3 μm step size of *z*-stacks using a 63× oil objective plus 4× digital zoom, then processed with the maximum-intensity projection using ImageJ. Quantification sample size: for filopodia numbers per MFT, 20-30 MFTs were collected per mouse; for PV puncta, density was averaged from 18 images per mouse; for dendritic spine densities, 20-30 dendrites were collected per mouse (Guo et al., 2018; Twarkowski et al., 2022).

### Ex vivo electrophysiology

Mice were unilaterally injected with 0.4 μL lenti-ablim3-shRNA/shNT and 0.3 μL AAV5-CamKII-ChR2-eYFP into the dorsal DG, and 0.3 μL AAV-S5E2-tdTomato into dorsal CA3/CA2 of 8-week-old mice (see above for stereotactic coordinates). At 2-3 weeks after viral infusion, mice were anaesthetized with ketamine and xylazine (10 mg/ml and 1.6 mg/ml, IP) then transcardially perfused with ice-cold (4 °C) choline chloride-based artificial cerebrospinal fluid (ACSF) composed of (in mM): 92 choline chloride, 2.5 KCl, 1.25 NaH2PO4, 30 NaHCO3, 20 HEPES, 25 glucose, and 10 MgSO4·7H2O. Their brains were rapidly extracted following decapitation. Coronal slices (300 μm thick) containing the dorsal hippocampus were cut in ice-cold (4 °C) choline chloride ACSF using a Leica VT1000 vibratome (Leica Biosystems) and transferred to warm (33 °C) normal ACSF for 30 min. Normal ACSF contained (in mM): 124 NaCl, 2.5 KCl, 1.25 NaH2PO4, 24 NaHCO3, 5 HEPES, 12.5 glucose, 2 MgSO4·7H2O, 2 CaCl2·2H2O. All ACSF solutions were adjusted to a pH of 7.4, mOsm of 305, and were saturated with carbogen (95% O2 and 95% CO2). Slices were allowed to cool to room temperature (20-22 °C) for 1 hour before recordings.

#### For whole-cell patch-clamp recordings

were obtained using a Multiclamp 700B amplifier (Molecular Devices) low-pass filtered at 1.8 kHz with a four-pole Bessel filter and digitized with a Digidata 1550B (Molecular Devices). Slices were placed in a submersion chamber and continually perfused (>2 mL/min) with normal ACSF. Neurons were visually identified by infrared differential interference contrast imaging combined with epifluorescence using LED illumination from a pE-300white (CoolLED). Pyramidal neurons in CA2 and CA3ab were distinguished by their anatomical location and distinct electrophysiological properties (Chevaleyre and Siegelbaum, 2010). Borosilicate patch pipettes had an impedance of 4-5 MΩ and filled with an internal solution containing (in mM): 120 CsMeS, 4 MgCl2, 1 EGTA, 10 HEPES, 5 QX-314, 0.4 Na3GTP, 4 MgATP, 10 phosphocreatine, 2.6 biocytin, pH 7.3, 290 mOsm. For miniature inhibitory postsynaptic current recordings, patch pipettes were filled with 140 mM CsCl in place of CsMeS. For current-clamp recordings, patch pipettes were filled with 130 mM potassium gluconate in place of CsMeS with QX-314 excluded. Once GΩ seal was obtained, neurons were held in voltage-clamp configuration at −70 mV and the input resistance, resting membrane potential, and capacitance were measured. Series resistance (<30 MΩ) was monitored throughout recordings and recordings were discarded if series resistance changed by >20% from baseline.

#### For electrically evoked EPSCs and IPSCs from DG

a concentric bipolar tungsten electrode was used to deliver electrical stimulation (0.2 ms duration) in the molecular layer of the DG. Electrically-evoked current responses were recorded at 1.5-fold threshold, defined as the minimum stimulation intensity required to produce a consistent current response beyond baseline noise.

#### For optically evoked EPSCs and IPSCs

2 ms 473 light pulses were delivered above the mossy fiber pathway – the hilus of the DG. Current responses were recorded at 1.5-fold threshold.

#### For current clamp recordings of PV INs

bursting neurons, identified by their asynchronous rapid firing within the first 50 ms of the current step followed by failure to fire, were excluded from analysis. For miniature postsynaptic current recordings, 1 μM tetrodotoxin was included in the bath ACSF. Kynurenic acid 2mM was included when recording mIPSCs and 0.1 mM picrotoxin when recording mEPSCs. Autodetection parameters for inclusion of events was determined by calculating minimum threshold: root mean square (RMS)2 × 1.5. To control for oversampling and unequal sample size between groups, quantile sampling of the frequency and amplitude was calculated by computing 29 evenly spaced quantile values from each neuron, starting at 1 percentile and ending at 94.2 percentile, with a step size of 3.33% (Hanes et al., 2020). Data acquisition was performed using Clampex and analyzed with Clampfit (Molecular Devices). EasyElectrophysiology software (V2.4.1) was used to analyze mPSC recordings.

### Statistics, rigor, and reproducibility

We adhered to accepted standards for rigorous study design and reporting to maximize the reproducibility and translational potential of our findings as described in (Landis et al., 2012)and in ARRIVE Guidelines (Kilkenny et al., 2010). All experimenters were blind to treatment conditions throughout data collection, scoring, and analyses. Statistical analyses were conducted using Prism v9 (GraphPad) and the minimum sample size was determined based on prior experience, existing literature, and a power analysis. Statistical significance was defined as p < 0.05. Two-tailed Student’s t-tests were used for two-group comparisons. Analysis of variance (ANOVA) were used for three or more group comparisons. Repeated-measures ANOVA were used for comparison of groups across treatment condition or time. Appropriate nonparametric tests were used when data sets failed to meet parametric assumptions. Detailed statistical analyses can be found in **Supplementary Table 1**.

## AUTHOR CONTRIBUTIONS

Conceptualization, AS; Methodology, JBA, YS, AS; Formal Analysis, JBA and YS; Investigation, JBA and YS; Resources, JBA, YS and AS; Writing, JBA, YS and AS; Visualization, JBA, YS and AS; Funding acquisition, AS

## ACKNOWLEDGEMENTS

We thank members of Sahay lab, L.M.S. Sahay and L.B.B. Sahay for comments on the manuscript. Y.S. was supported by MGH ECOR Fund for Medical Discovery (FMD) Fundamental Research Fellowship Award and is recipient of a Harvard Brain Initiative Travel Grant. A.S. acknowledges support from NIH 1R01MH111729-01, 3R01MH111729-04S1, R01 AG076612-02, 3R01AG076612-01S1 diversity supplement, The Simons Collaboration on Plasticity and the Aging Brain, James and Audrey Foster MGH Research Scholar Award and MGH department of Psychiatry.

## DECLARATION OF INTERESTS

The authors declare no competing interests.

## KEY RESOURCES TABLE

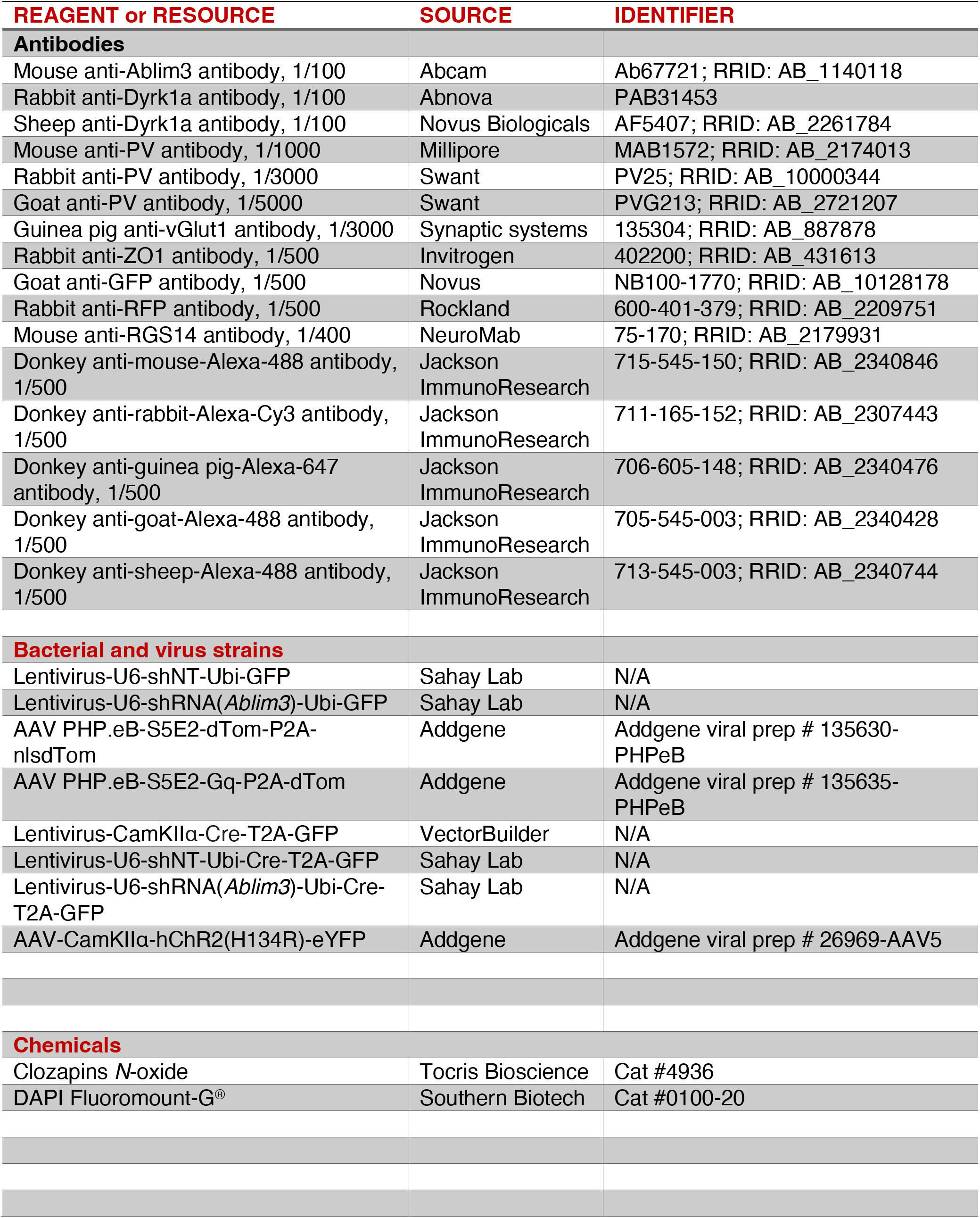
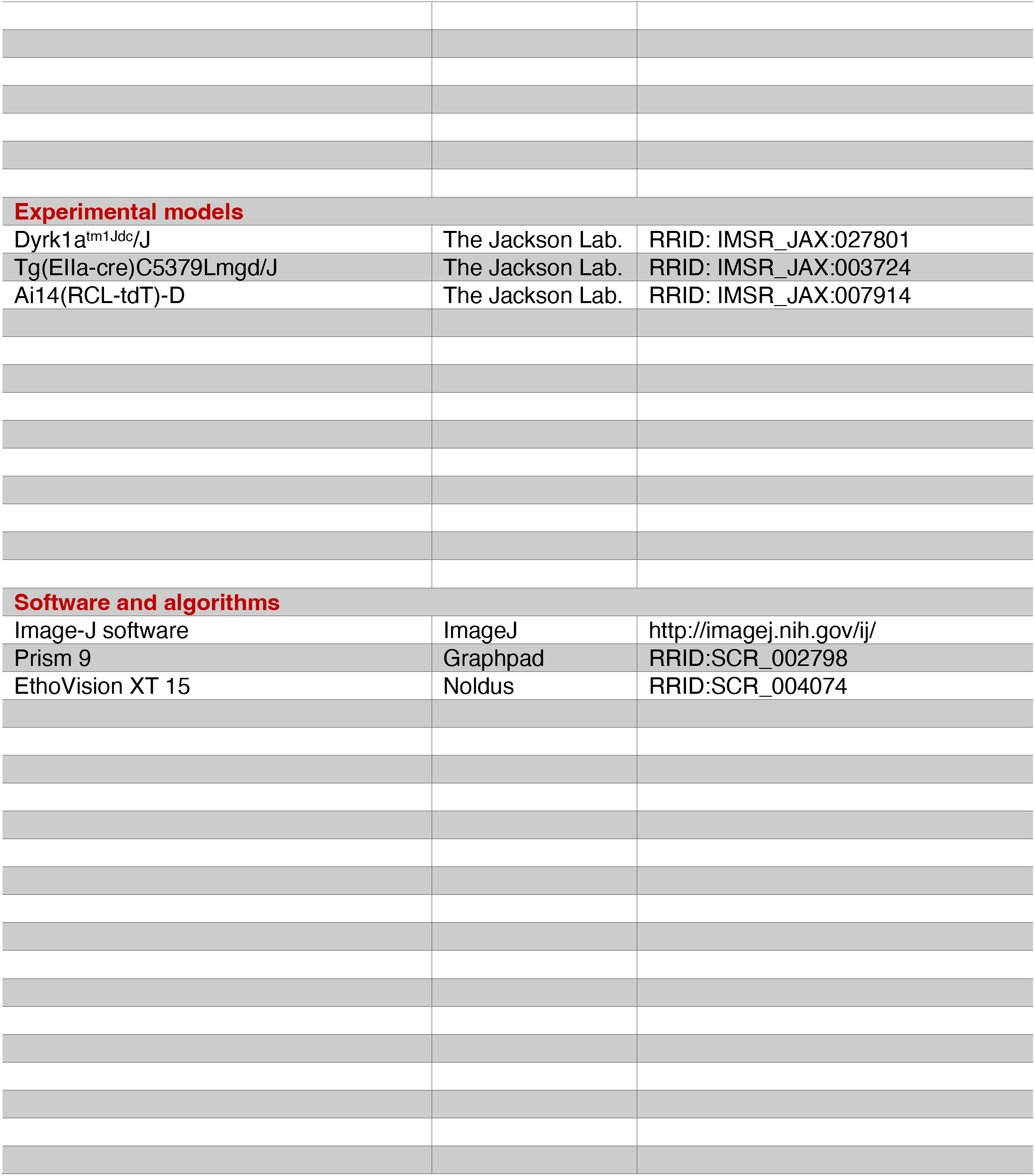

## Supplementary Figures

**Figure S1.**
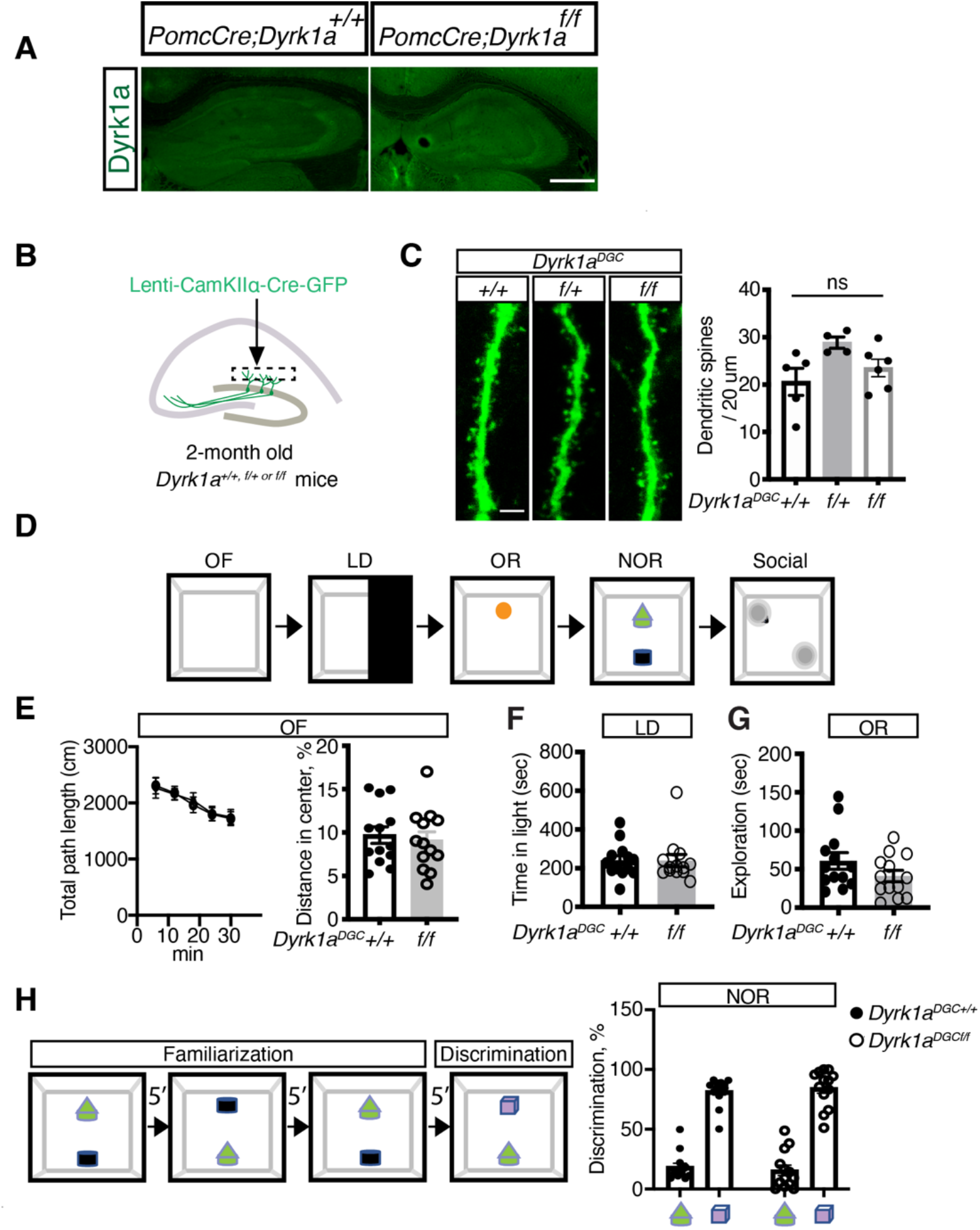
Loss of *Dyrk1a* in DGCs does not affect dendritic spine density, locomotion, anxiety-like behavior and novel object recognition. **Related to Figure 1.** (A) Dyrk1a Immunostaining in hippocampus of *PomcCre;Dyrk1a^+/+^* and *PomcCre:Dyrkla^f/f^* mouse. Note the complete loss of signal in DG of *PomcCre:Dyrkla^f/f^* brain section (mossy fibers are not labeled) indicating specificity of Dyrk1a antibody. Scale bar, 500 μm. (B) Schematic of injection of lenti-CamKIIα-Cre-T2A-GFP viruses into DG of 2-month old *Dyrk1a^+/+, f/+ or f/f^* mice 2 weeks prior to behavioral testing. (C) Representative images showing GFP^+^ dendrites and quantification of dendritic spine density (N = 5 for mice +/+; 4 mice for f/+; 6 mice for f/f) in outer one-third of the molecular layer of the DG. Scale bar, 2 μm. ns, not significant using one-way ANOVA with Bonferroni post hoc test. (D) Schematic of behavior testing schedule. OF, open-field task; LD, light-dark task; OR, object recognition; NOR, novel object recognition; Social cognition task (see Figure 1H). (E) OF, quantification of total distance travelled (left) and percentage of distance traveled across the center arena (right). (F) LD box assay, quantification of the time spent in the light compartment. (G) OR, object recognition. Quantification of the time spent sniffing the object (seconds). (H) NOR, novel object recognition, mouse was exposed to 2 objects over three sessions (5 minutes each session, object locations counterbalanced) followed by a fourth session in which one object was replaced with a new object. Time spent sniffing the objects was quantified. All data are displayed as mean ± SEM and analyzed using two-tailed, unpaired *t* test with Welch’s correction (E-G) or two-way ANOVA with Bonferroni post hoc test (H).

**Figure S2.**
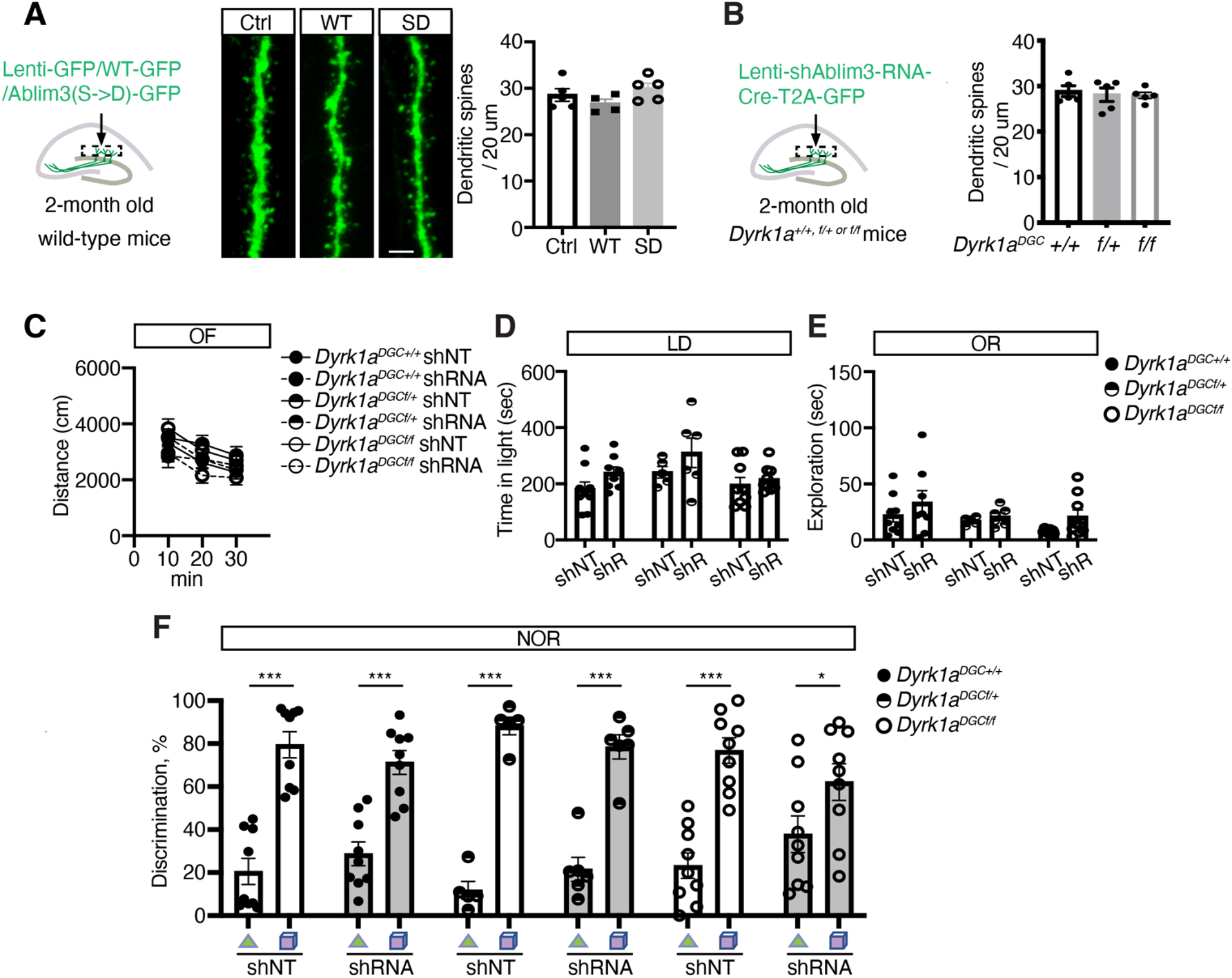
Viral manipulations of *Ablim3* in DGCs of wild-type or *Dyrk1a^DGC f/+ or f/f^* mice and characterization of dendritic spine density, locomotion, anxiety-like behavior and novel object recognition. **Related to Figure 2.** (A) Representative images showing GFP^+^ dendrites and quantification of dendritic spine density in the outer one-third of the molecular layer of the DG. Scale bar, 2 μm. ns, not significant using one-way ANOVA with Bonferroni post hoc test. (B) Schematic of lenti-shRNA (*Ablim3*)-Ubi-Cre-T2A-GFP viruses injection into DG of 2-month old mice for 2 weeks (left) and quantification of dendritic spine densities (right). ns, not significant using one-way ANOVA with Bonferroni post hoc test. (C) OF, quantification of total distance travelled (left) and percentage of distance traveled across the center arena (right). (D) LD box assay, quantification of the time spent in the light compartment. (E) OR, object recognition. Quantification of the time spent sniffing the object (seconds). (F) NOR, novel object recognition, mouse was exposed to 2 objects over three sessions (5 minutes each session, object locations counterbalanced) followed by a fourth session in which one object was replaced with a new object. Time spent investigating/sniffing the objects was quantified. All data are displayed as mean ± SEM and analyzed using two-way ANOVA with Bonferroni post hoc test (D-F).

**Figure S3.**
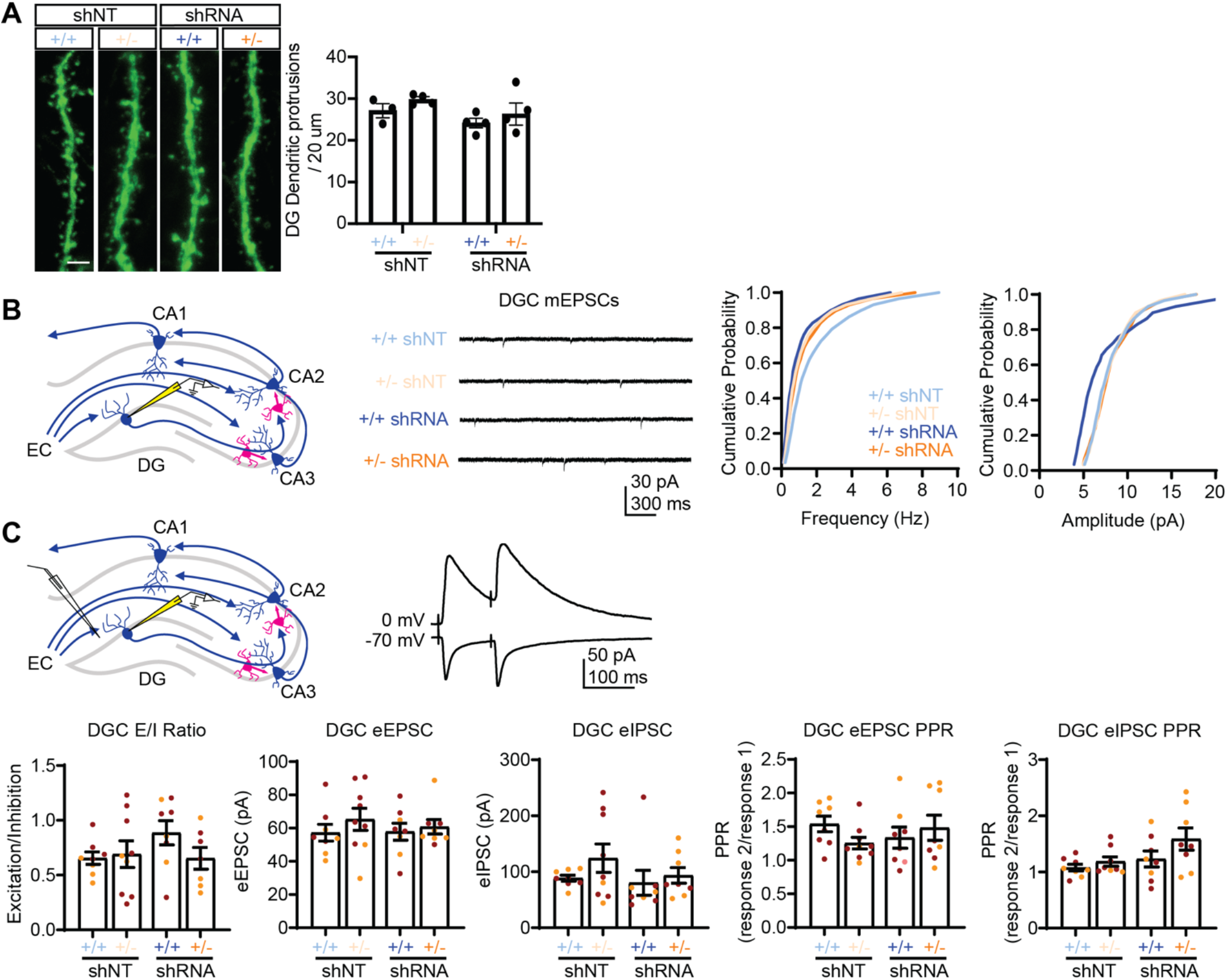
Characterization of perforant path-DGC circuitry in *Dyrk1a* hemizygous mice. Related to Figure 3. (A) Representative images showing GFP^+^ dendrites and quantification of dendritic spine density in the outer one-third of the molecular layer of the DG. Scale bar, 2μm. *Dyrk1a^+/+^* mice: N = 3 mice for shNT; N=4 mice for shRNA; in *Dyrk1a^+/-^* mice: N=4 mice for each group. Not significant using two-way ANOVA with Bonferroni post hoc test. (B) Schematic depicting whole-cell patch-clamp recording of mEPSC from DGCs (left). Representative recording trace from each group (middle). Cumulative probability plots of mEPSC frequency and amplitude from DGCs (Kolmogorov-Smirnov test, *\*p* < 0.05, n=8-9 cells, 2-3 cells per mouse, 4-5 mice per group). (C) Schematic depicting whole-cell patch-clamp recording of EPSC and IPSC from DGC to paired pulse electrical stimulation delivered onto the molecular layer of the dentate gyrus. Representative trace depicting EPSC and IPSC from a DGC. Bar graphs indicate excitation to inhibition ratio, the amplitude of the first EPSC and IPSC response to paired pulse electrical stimulation, and the EPSC and IPSC paired pulse ratios. (Two-way ANOVA with Tukey posthoc, \**p* < 0.05, n=8-9 cells, 2-3 cells per mouse, 3-4 mice per group).

**Figure S4.**
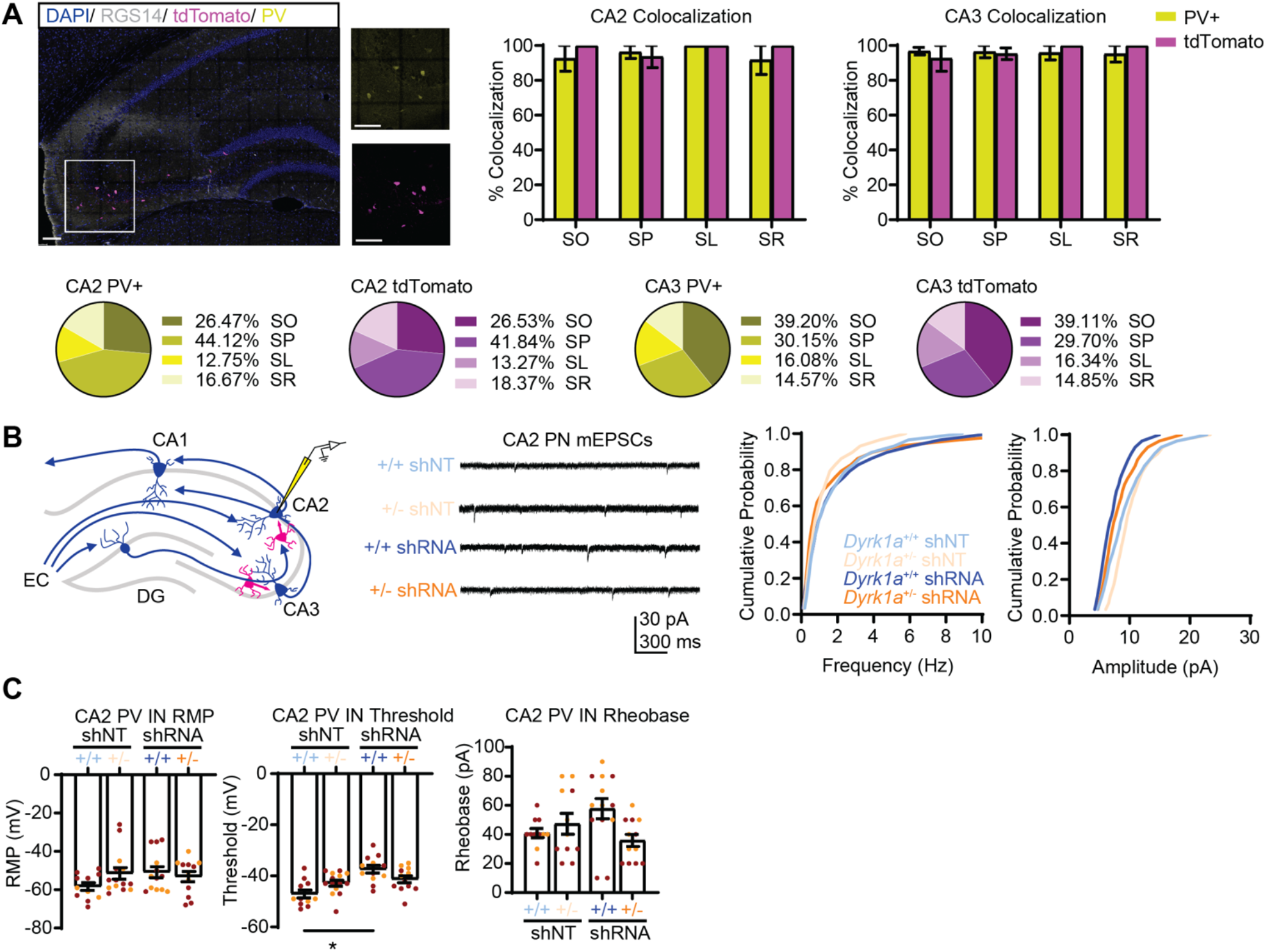
Characterization of excitatory synaptic inputs onto CA2 PN and CA2 PV IN passive membrane properties in adult *Dyrk1a^f/+^* mice and following *Ablim3* downregulation in DGCs. **Related to Figure 4.** (A) Representative images depicting expression of DAPI, AAV-S5E2-tdTomato, and PV immunostaining congruency, with RGS14 immunostaining of CA2. Scale bar, 100 μm. Graphs indicate percent colocalization of S5E2-tdTomato+ and PV+ neurons across CA2 and CA3 hippocampal layers: (SO) stratum oriens, (SP) stratum pyramidale, (SL) stratum lucidum, (SR) stratum radiatum. Not significant using two-way ANOVA with Bonferroni post hoc test. Pie charts indicate distribution of S5E2-tdTomato^+^ and PV^+^ neurons across CA2 and CA3 hippocampal layers. (B) Schematic depicting whole-cell patch-clamp recording of mEPSC from CA2 pyramidal neuron (left). Representative recording trace from each group (middle). Cumulative probability plots of mEPSC frequency and amplitude from PV INs (Kolmogorov-Smirnov test, *\*p* < 0.05, n=8-9 cells, 2-3 cells per mouse, 4-5 mice per group). (C) Bar graphs of CA2 PV IN passive membrane properties (Two-way ANOVA with Tukey posthoc, \**p* < 0.05, n=12-14 cells, 2-3 cells per mouse, 5-6 mice per group).

**Figure S5.**
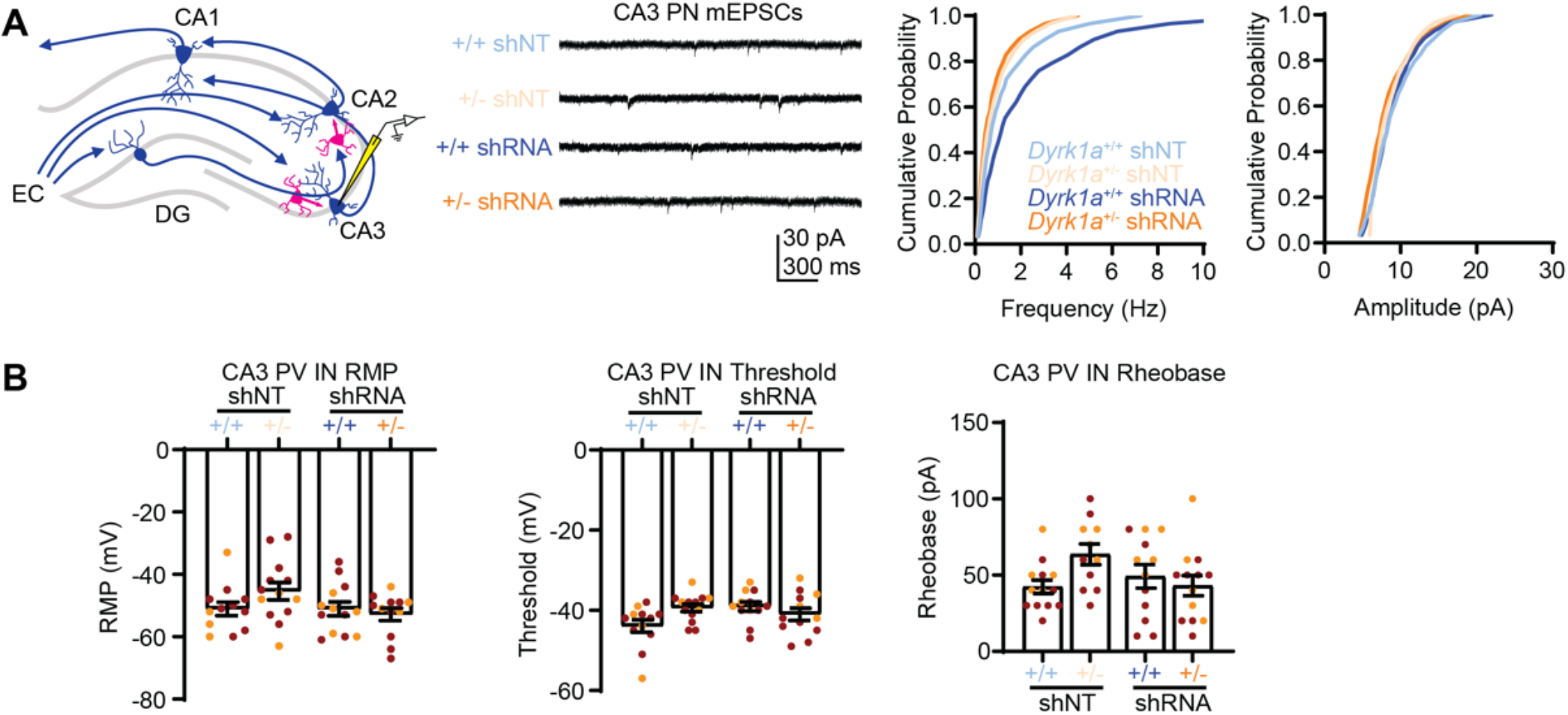
Characterization of excitatory synaptic inputs onto CA3 PN and CA3 PV IN passive membrane properties in adult *Dyrk1a^f/+^* mice and following Ablim3 downregulation in DGCs. **Related to Figure 5.** (A) Schematic depicting whole-cell patch-clamp recording of mEPSC from CA3 pyramidal neuron (left). Representative recording trace from each group (middle). Cumulative probability plots of mEPSC frequency and amplitude (Kolmogorov-Smirnov test, \**p* < 0.05, n=8-9 cells, 2-3 cells per mouse, 4-5 mice per group). (B) Bar graphs of CA3 PV IN passive membrane properties (Two-way ANOVA with Tukey posthoc, \**p* < 0.05, n=12-13 cells, 2-3 cells per mouse, 5-6 mice per group).

**Figure S6.**
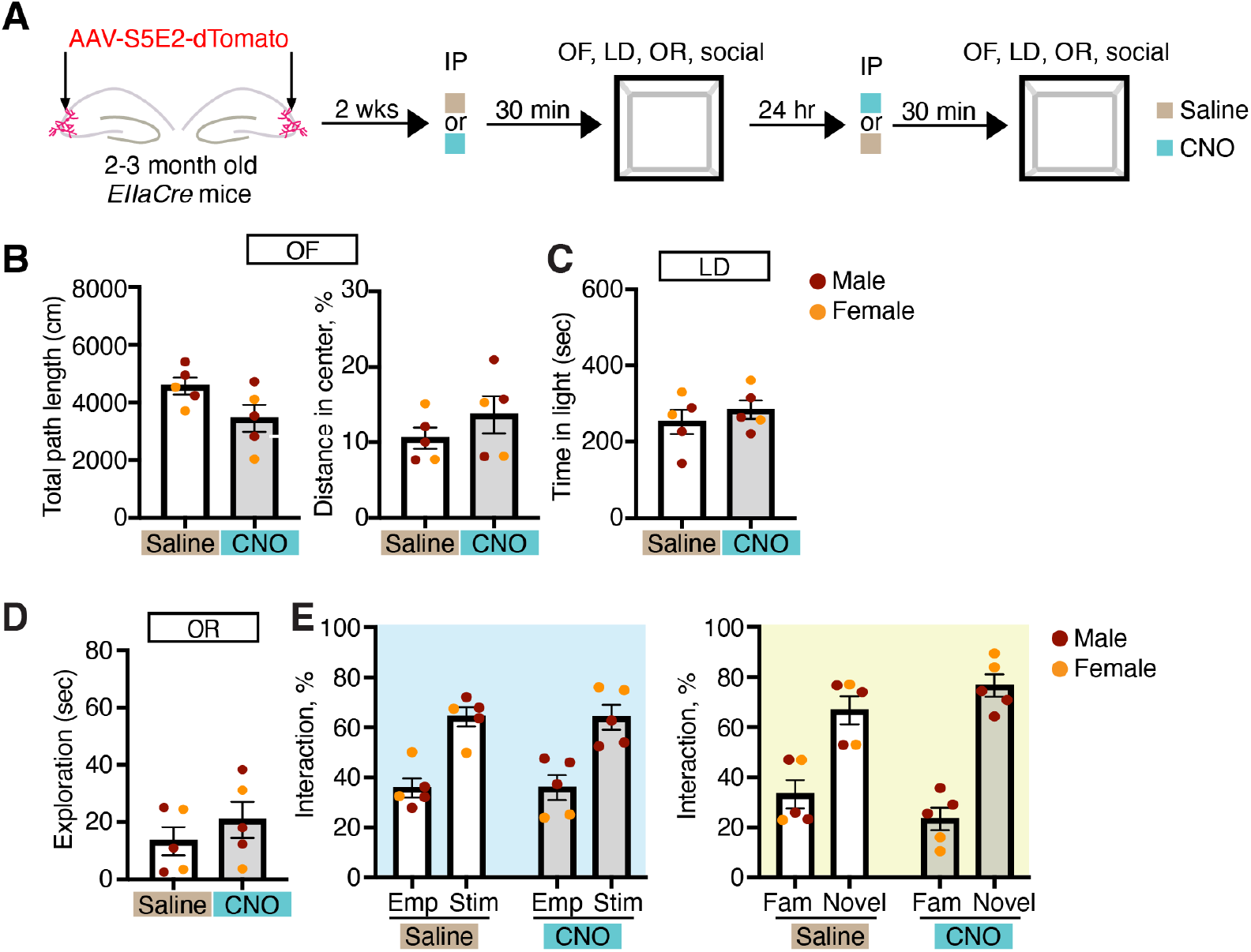
CNO treatment does not affect anxiety-like, object recognition or social recognition. **Related to Figure 7.** (A) Schematic of virus injection and behavioral testing timeline. (B) OF, quantification of total distance travelled (left) and percentage of distance traveled across the center arena (right). (C) LD box assay, quantification of the time spent in the light compartment. (D) OR, object recognition. Quantification of the time spent sniffing the object (seconds). (E) Quantification of empty cup vs. social stimulus interaction time during social recognition phase (T1) (left panel, blue shaded) and the discrimination trial (T4, right panel, yellow shaded). N= 5 mice for each group. All data are displayed as mean ± SEM. and analyzed using two-way ANOVA with Bonferroni post hoc test.

## Notes

### Competing Interest Statement

The authors have declared no competing interest.

